# A small molecule enhances arrestin-3 binding to the β_2_-adrenergic receptor

**DOI:** 10.1101/2024.12.12.628161

**Authors:** Han Kurt, Ali Akyol, Cagdas Devrim Son, Chen Zheng, Irene Gado, Massimiliano Meli, Erica Elisa Ferrandi, Ivan Bassanini, Francesca Vasile, Vsevolod V. Gurevich, Aylin Nebol, Esra Cagavi, Giulia Morra, Ozge Sensoy

**Author notes:** Corresponding authors: Giulia Morra and Ozge Sensoy. **Email:**. ^Ŧ^ Contributed equally to this work (Han Kurt, Ali Akyol). **Author Contributions:** GM and OS designed the research. OS funding acquisition. HK performed MD simulations and virtual screening. HK, GM, and OS analyzed data. MM assisted in virtual screening. AA performed FRET experiments. CZ performed NanoBiT experiments, CZ and VVG analyzed and interpreted NanoBiT data. AA and CDS analyzed and interpreted FRET data. AN performed experiments with cardiac cell culture. AN and EC analyzed and interpreted data. EF, IB, IG and FV performed NMR experiments. IG and FV interpreted NMR data. HK, AA, AN, CDS, EC, FV, GM, VVG, and OS contributed to writing the paper.

## Abstract

G protein-coupled receptor (GPCR) signaling is terminated by arrestin binding to a phosphorylated receptor. Binding propensity has been shown to be modulated by stabilizing the pre-activated state of arrestin through point mutations or C-tail truncation. Here, we hypothesize that pre-activated rotated states can be stabilized by small molecules, and this can promote binding to phosphorylation-deficient receptors, which underly a variety of human disorders. We performed virtual screening on druggable pockets identified on pre-activated conformations in Molecular Dynamics trajectories of arrestin-3, and found a compound targeting an activation switch, the back loop at the inter-domain interface. According to our model, consistent with available biochemical and structural data, the compound destabilized the ionic lock between the finger and the back loop, and enabled transition of the ‘gate loop’ towards the pre-activated state, which stabilizes pre-activated inter-domain rotation. The predicted binding pocket is consistent with saturation-transfer difference NMR data indicating close contact between the piperazine moiety of the compound and C/finger loops. The compound increases in-cell arrestin-3 binding to phosphorylation-deficient and wild-type β2-adrenergic receptor, but not to muscarinic M2 receptor, as verified by FRET and NanoBiT. This study demonstrates that the back loop can be targeted to modulate interaction of arrestin with phosphorylation-deficient GPCRs in a receptor-specific manner.

## Introduction

G protein-coupled receptors (GPCRs) are involved in various physiological and pathophysiological processes and targeted by about a third of clinically used drugs ^1^. Agonist binding to GPCRs triggers a series of conformational rearrangements, which are propagated to the cytoplasmic tip of the receptor^2^. Active GPCR conformation initiates G protein-mediated signaling. G protein-coupled receptor kinases (GRKs) phosphorylate certain serine and threonine residues on the cytoplasmic regions of the activated receptor^3^. Consequently, active and phosphorylated receptor recruits another cytosolic protein, arrestin, which terminates G protein-mediated signaling and ^4–6^ activates alternative signaling pathways ^7–9^.

Mammals express four arrestin subtypes: arrestin-1 and arrestin-4 are known as visual arrestins and expressed in photoreceptors, whereas arrestin-2 (β-arrestin-1) and arrestin-3 (β-arrestin-2) are known as non-visual arrestins and expressed in virtually every cell in the body^*^. All arrestins are elongated molecules consisting of an N- and C-domain^10^. In the basal state, the carboxy-terminal tail (C-tail) of arrestin folds back and is anchored to the N-domain of the protein, thereby stabilizing the basal conformation. Arrestin binding to the GPCR requires receptor attached phosphates and an active receptor conformation^11–14^. Arrestin protein family members show different propensity towards the phosphorylation state of the receptor. For instance, arrestin-1 and arrestin-4 require receptor phosphorylation for binding to rhodopsin while requirement of non-visual arrestin-2 and arrestin-3 for phosphorylation depends on the receptor type^3^.

Receptor binding induces several conformational rearrangements in arrestin, the most prominent being the release of the C-tail and twisting of the two domains relative to each other by ∼20° ^15–21^. These are triggered by the phosphorylated tail of the receptor that binds to the N-domain of arrestin, displacing the C-tail and releasing the structural constraints that stabilize its basal conformation. The interdomain rotation exposes the interface regions engaged in the formation of the high-affinity complex with the receptor.

To understand the determinants of pre-activation of arrestin, various sequence modifications have been tested, such as truncation of C-tail and point mutations. These attempts resulted in experimentally solved structures with different inter-domain rotation angles (7.5° in the pre-‘polar cor’ arrestin-1 mutant^22^; ∼20 ° in the splice variant of visual arrestin^23^) suggesting a different impact on activation. The effect of PIP2 binding has also recently been probed via HDX exchange^24^ confirming the allosteric communication between N- and C-domain and vice versa^25^ underlying activation.

Pre-activated constructs with destabilized basal conformation can bind non-phosphorylated GPCR. In that respect, they can be considered as potentially useful tools for binding phosphorylation deficient receptors which underlies a variety of human disorders, including *retinitis pigmentosa*^26^, certain forms of cancer^27^, diabetes ^28^, and others^29^. Nevertheless, such constructs usually demonstrate reduced thermal stability and receptor selectivity^30^.

An alternative approach for enhancement of interaction between arrestin and non-phosphorylated or phosphorylation-deficient GPCR is the stabilization of pre-activated arrestin conformations by small molecules. Arrestin-3 is intrinsically flexible enough and can transiently sample pre-activated conformations, as shown in a computational study^31^. The pre-activated states are characterized by an exposed C loop (residues 249-254 in arrestin-2 and 244-249 in arrestin-3), a rearranged gate loop (residues 295-306 in arrestin-2 and 290-299 in arrestin-3)^32^ inter-domain rotation, and unfolded back loop (residues 313-317 in arrestin-3, 312-316 in arrestin-2; 318-322 in arrestin-1) ^23,33,34^. Here, we tested whether stabilization of such pre-activated states enhances arrestin-3 recruitment to a non-phosphorylated GPCR. A computational drug discovery approach, based on targeting pre-activated arrestin-3 conformations extracted from MD simulations, identified a small molecule which was predicted to bind arrestin-3. The binding of the molecule was verified by saturation-transfer difference (STD) NMR. We verified that the compound enhanced arrestin-3 binding to phosphorylation-deficient and wild-type β2-adrenergic receptor, but not to muscarinic M2 receptor, by FRET and NanoBiT, demonstrating receptor specificity.

## Results

### Free arrestin-3 samples pre-activated conformations

Molecular Dynamics (MD) simulations of free arrestin-3 revealed a marked flexibility of the interdomain interface ^33^. To explore spontaneously adopted pre-activated states for *in silico* drug discovery, we ran three MD simulations, each of which was 1 μs, in explicit water. The relation between the conformational state of the back loop, inter-domain rotation, and the rearrangement associated with the gate loop occurring upon GPCR binding^15–21,35^, motivated us to use them as reaction coordinates to identify pre-activated states of arrestin-3. We focused on the distances between the Cα atoms of residues K313 and G317 located at the termini of the back loop (blue in Figure 1A) in the inactive state, and residues D291 and H296 on the gate loop (red in Figure 1A), which report on conformational changes occurring upon receptor binding. We classified conformations extracted from simulations as inactive/pre-activated/active (see Methods for definitions and Figure 1B for distributions) based on the reference values in the corresponding structures. Arrestin-3 sampled inactive/pre-activated/active states of the back loop in (68/29/3)%, (59/40/1)%, and (15/54/31)% of the three independent simulations. Also, pre-activated, albeit not fully active, states of the gate loop were sampled in 72%, 60%, and 50% of them, respectively. The population of active and pre-activated states was confirmed in a separate set of accelerated molecular dynamics (aMD) simulations both for back loop, with a distribution of inactive/pre-activated/active states (61/33/6)%, respectively, and gate loop (pre-activated state sampled in 78% of the MD trajectory). We also analyzed the status of the aromatic core at the interdomain interface, whose destabilization was proposed as an activation-related change^33^. It adapted inactive/pre-activated/active states in (10/79/11)%, (12/41/46)% and (17/82/1)% of the three arrestin-3 trajectories. However, we did not consider targeting this region as it is involved in GPCR binding, interacting with the intracellular loop 2 of the receptor^15^ .

**Figure 1.**
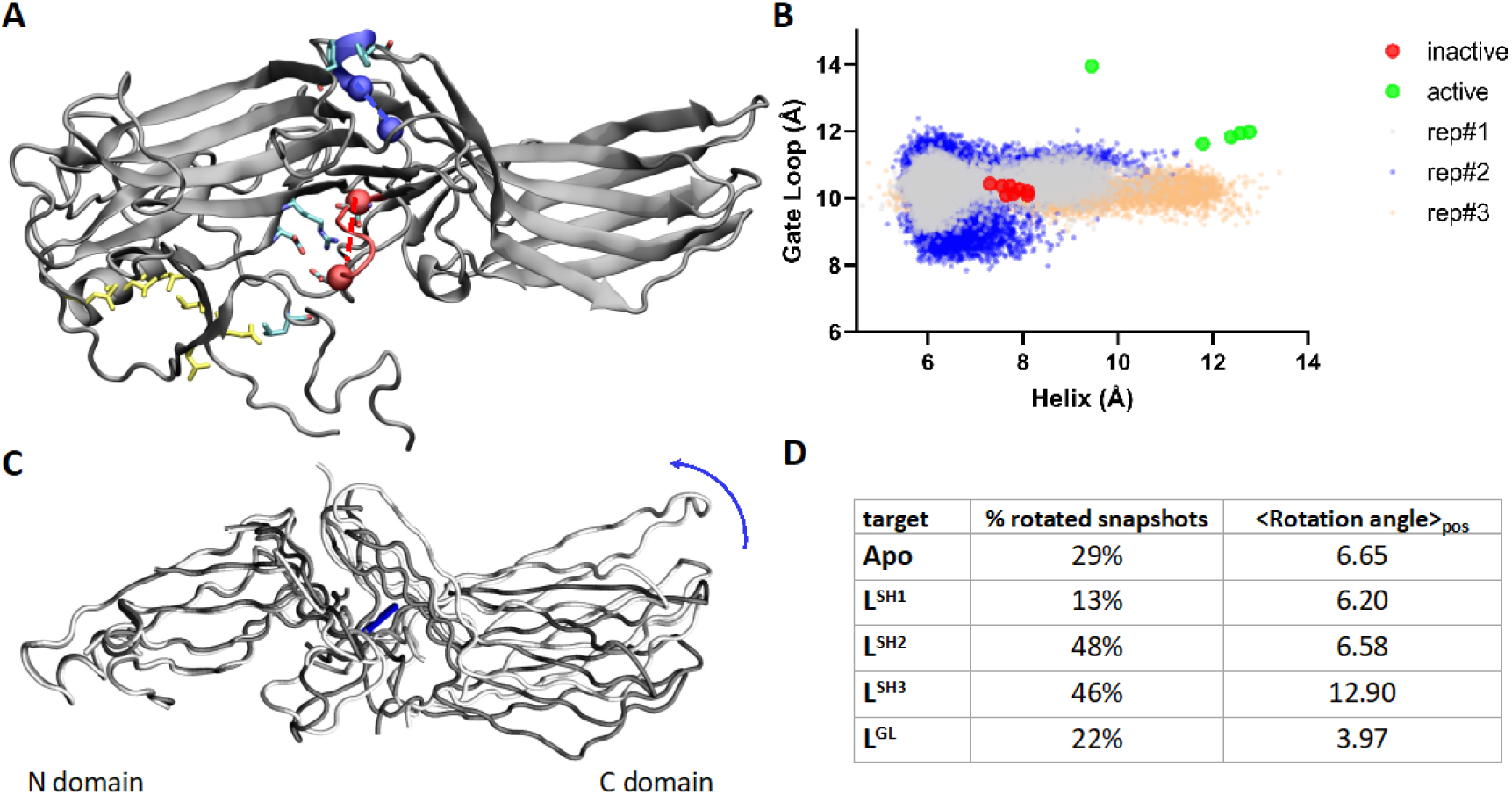
**A**. Distances between residues K313 and G317 at the termini of the back loop *(blue),* and residues D291 and H296 for gate loop (red) are shown on the 3D structure of arrestin-3 (PDB ID: 3P2D)^31^. **B**. Distribution of the distances used to identify the conformational state of the gate loop and the back loop in arrestin-3 trajectories and those measured in active/pre-activated and inactive crystal structures of arrestins (See Methods for the complete list of the crystal structures). Values for inactive and active crystal structures are shown in red and green dots, respectively. Notably, inositol hexakisphosphate-bound arrestin-3 (PDB ID:5TV1) ^16^ adopts an extremely long distance (See green dot outlier) at the gate loop, due to the perturbation induced by the IP6 molecule near the polar core. **C** Superposition of traces of two extreme conformations from the principal component analysis of arrestin-3 crystal structures, aligned on the N terminal domain, showing the rotation axis and direction (See Methods). **D** Table summarizing the occurrence of rotated states in arrestin-3-compound complexes and the average rotation angle.

### Targeting the back loop in the unfolded state stabilizes the pre-activated inter-domain rotation in arrestin-3

Starting from the pre-activated states sampled in MD simulations we set out to find small molecules that stabilize these conformations. Pre-activated conformational states were sufficiently populated in the MD trajectory dataset to extract selected snapshots for *in silico* drug discovery. We chose seven structures characterized by a diverse set of conformational states of the back loop and the gate loop as well as the interdomain rotation (See Methods for the description of the conformational states, and Supplementary Fig.1 and Supplementary Table 1 for associated values of the back loop, gate loop and inter-domain rotation). We then searched for available binding pockets on the interdomain interface using these structures and found two pockets with the best druggability scores (Supplementary Table 2), one of which was located at the back loop and the other at the gate loop. Virtual screening of these pockets yielded candidate compounds, which were further analyzed by MD simulations of the predicted complexes with arrestin-3. One compound, referred to as L^GL^, bound stably to the gate loop throughout the course of 1 μs simulation. On the back loop pocket, we selected 10 candidate compounds (L^SH^ series) based on the docking scores (Supplementary Table 3 and Supplementary Fig.2 for examples to binding poses) which stably bound to the back loop in 1-μs MD trajectory.

We measured the predicted inter-domain rotation for the stable complexes in the MD simulations. Within the L^SH^ series (Fig. 1D), we consistently observed the perturbation of the rotation angle for L^SH-3^ (See Fig.2B, C for the predicted binding sites of the compounds). This complex adapts a rotation angle in 46% of the trajectories with an average value of 12.9^0^. L^SH-1^ and L^SH-2^, as well as L^GL^ (See Fig2.D for the predicted binding site of the compound), stabilized a lower rotation angle ∼6.5^0^ (Fig. 1D). To test whether compounds stabilize the binding competent conformation of arrestin-3, we also examined the dynamics of the back loop and the gate loop in the arrestin-3/compound complexes. The gate loop parameters (RMSD and the distance between Cα atoms of D291 and H296, both of which were expected to increase upon activation), are more perturbed with bound L^SH-3^ than in the other complexes, in agreement with the trend observed for the inter-domain rotation angle (See Figure 1D). Also, the distance between the gate loop and the polar core^25^, in arrestin-3 /compound trajectories, monitored by the side chains of D291 and R170 (D296 and R175 in arrestin-1) increases (5.0 Å) with L^SH-3^ relative to the crystal of inactive arrestin-3 (PDB ID: 3P2D).

**Figure 2.**
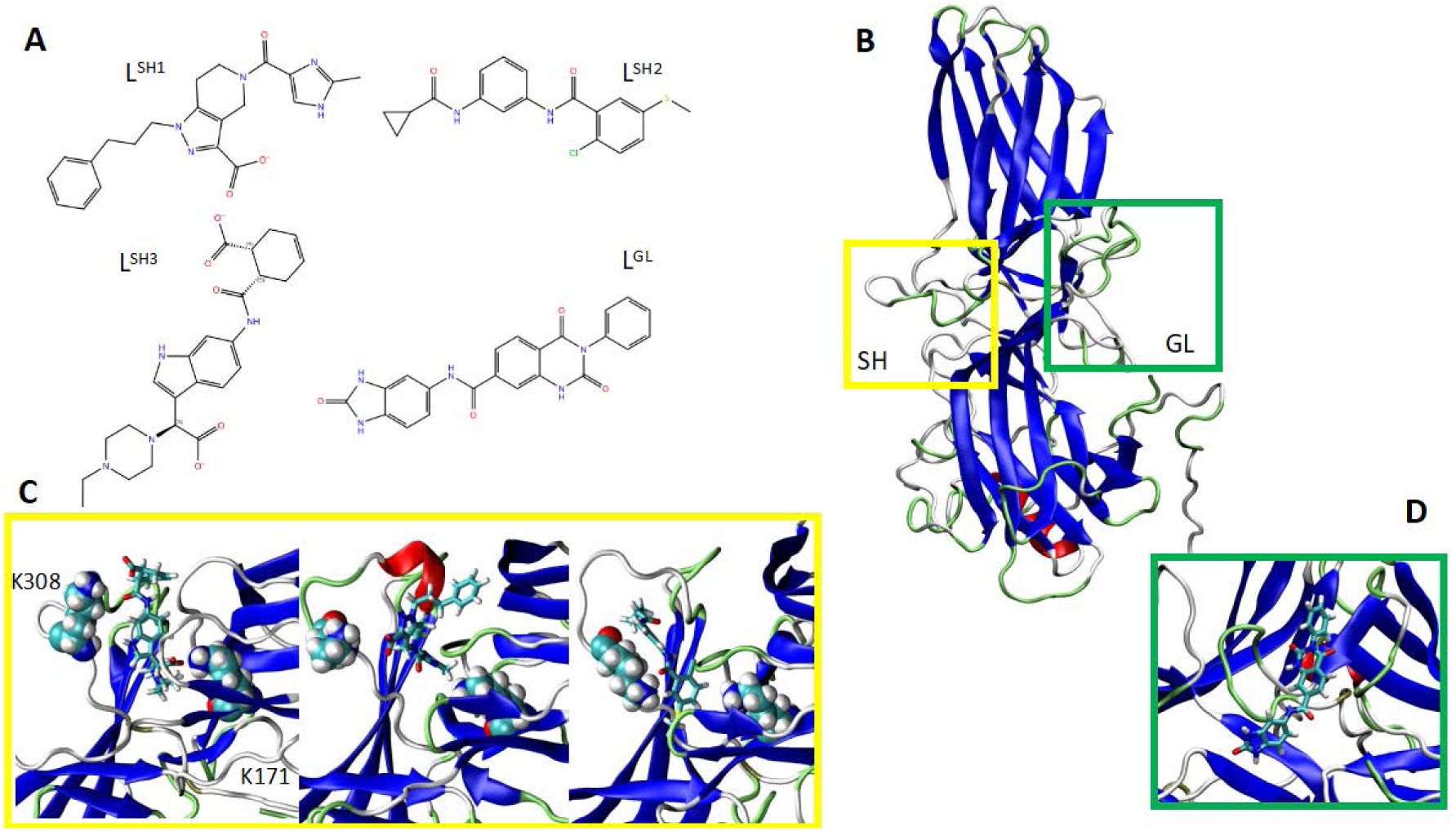
**A**. Structures of the compound candidates emerging from virtual screening. **B.** All atom structure of inactive arrestin-3 highlighting the two interface regions targeted by drug discovery, namely the back loop characterized by the short helix (SH) and the *gate loop* (GL). **C.** Close-up of the SH pocket with bound compounds: from top to bottom, L^SH-3^, L^SH-1^, L^SH-2^. **D.** Close-up of the GL pocket with bound compound L^GL^.

Interestingly, the perturbation of the back loop induced by the compounds is not always associated with the stabilization of pre-activated inter-domain rotation. For instance, L^SH-1^ stabilized the back loop in the unfolded state, but this did not perturb the gate loop, leading to sampling relatively lower inter-domain rotation (See Supplementary Fig.3). This suggests that both stabilization of the back loop in the unfolded state and perturbation of the gate loop towards pre-activated state are required to sample higher inter-domain rotation angles.

### L^SH-3^ binds arrestin-3 at the interdomain interface

To validate the interaction between arrestin-3 and L^SH-3^, Saturation Transfer Difference (STD) NMR was employed. STD-NMR is a powerful biophysical method for identifying binding between small molecules and larger molecules, facilitating the screening of compounds and their binding activity^36^. The STD NMR spectrum and the binding epitope (the protons in close contact with the protein) are shown in Fig. 3. Protons H14 and H15 of the piperazine ring showed the highest absolute STD%, suggesting that this moiety is the closest to the protein. A high STD value is also displayed by the methyl group (H17), while proton H16 could not be considered due to proximity of its signal to DMSO peak. Thus, piperazine ring and its short aliphatic chain effectively engage the binding pocket in arrestin-3. The aromatic (H7, H8 and H9) and cyclohexene (H1, H2, H3, H4 and H5) protons demonstrate lower intensity (0.3-1.5 STD%), indicating their weaker interactions with the protein surface. Thus, the molecule is arranged in the binding pocket of arrestin-3 with the piperazine moiety having the strongest interaction with the protein.

**Figure 3.**
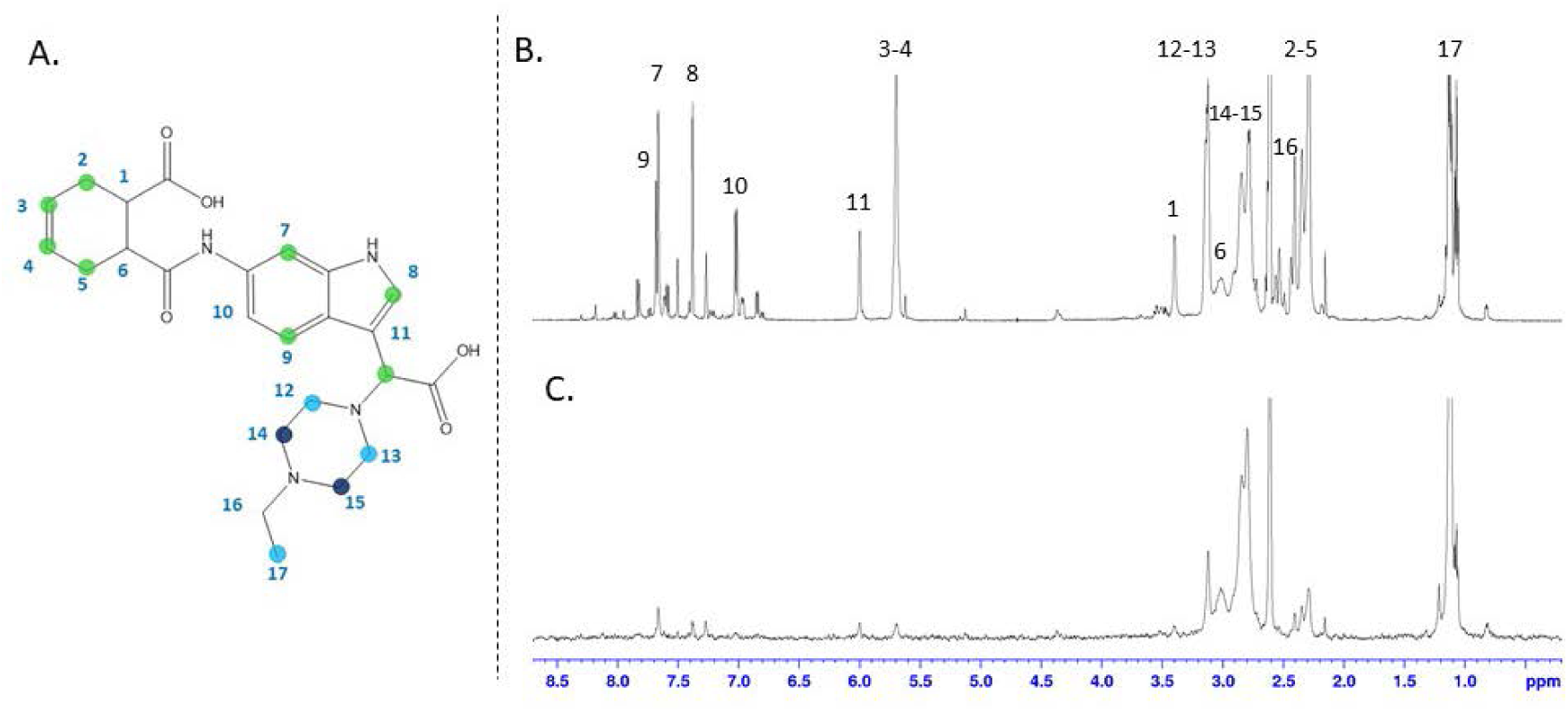
**A.** Epitope map of L^SH-3^ with relative STD percentages at initial slope conveyed by color code, acquired at saturation time of 2s, irradiating at 0 ppm. Dark blue dots indicate the most intense signal (100% relative STD), light blue dots between 20% and 80% and green dots under 20%. **B.** ^1^H spectrum of L^SH-3^ **C.** STD spectrum of L^SH-3^ in the presence of arrestin-3.

While STD experiments identify the functional groups of the compound critical for target binding, they do not identify the protein binding site. To evaluate which arrestin-3 site is involved, differential epitope mapping by STD NMR spectroscopy (DEEP-STD NMR)^37^ was employed. This approach requires the protein saturation at different irradiation frequencies to highlight parts of the compound contacting protein in the bound state. The analysis was carried out irradiating at 0 ppm and 8.8 ppm, centered on the methyl group of the aliphatic chains and the aromatic or NH protons, respectively. These two STD NMR experiments (Supplementary Fig.4) showed different compound epitopes under different irradiations, suggesting that the aromatic protons of the compound are more intense upon the irradiation of the protein at 8.8, while the piperazine ring is surrounded by aliphatic amino acids. The two main putative binding sites emerging from the computational site mapping, SH-short helix (from the helix turn present in the inactive back loop) and GL-gate loop (Fig. 2), differ in the amino acid composition: higher number of hydrophobic residues is present in the SH binding site, while charged residues are mainly present at GL (Supplementary Fig. 2). To discriminate among them, the total compound saturation was calculated for both STD spectra. A larger total saturation was observed in the STD spectrum irradiated at 0 ppm (Supplementary Table 5), indicating that the compound preferentially occupies the site lined by hydrophobic residues.

Finally, the binding constant (K_D_) of the compound was determined using titration STD experiments. The curves were built with four concentrations (from 0.3 mM to 5 mM) and the K_D_ values were determined from the binding isotherm with STD-AF measured at saturation times of 2s. (Supplementary Fig. 6). To calculate the STD Amplification Factor (STD-AF), which relates the protons’ proximity to the protein surface, protons H14, H15, and H17 were selected. The experimental data were fitted with an exponential function. The K_D_ obtained by STD-NMR is 0.8 mM, indicating that L^SH-3^ has sufficient affinity for arrestin-3 to be used as the chemical scaffold.

### L^SH-3^ enhances recruitment of arrestin-3 to phosphorylation-deficient β_2_AR in cells

β_2_AR C-tail contains four residues which can be phosphorylated by GRK2: Ser^355,356,364^ and Thr^360^. We replaced these residues with alanines to generate a phosphorylation-deficient mutant, as described^38^. We visualized arrestin-3 recruitment to wild-type (WT) and phosphorylation-deficient β_2_AR by fluorescent microscopy in response to an agonist isoproterenol (Iso) ^39^. Since this interaction requires the phosphorylation of the receptor, GRK2 was also added. Arrestin-3 was recruited to WT, but not phosphorylation-deficient β_2_AR (Fig. 4F and 4I).

**Figure 4.**
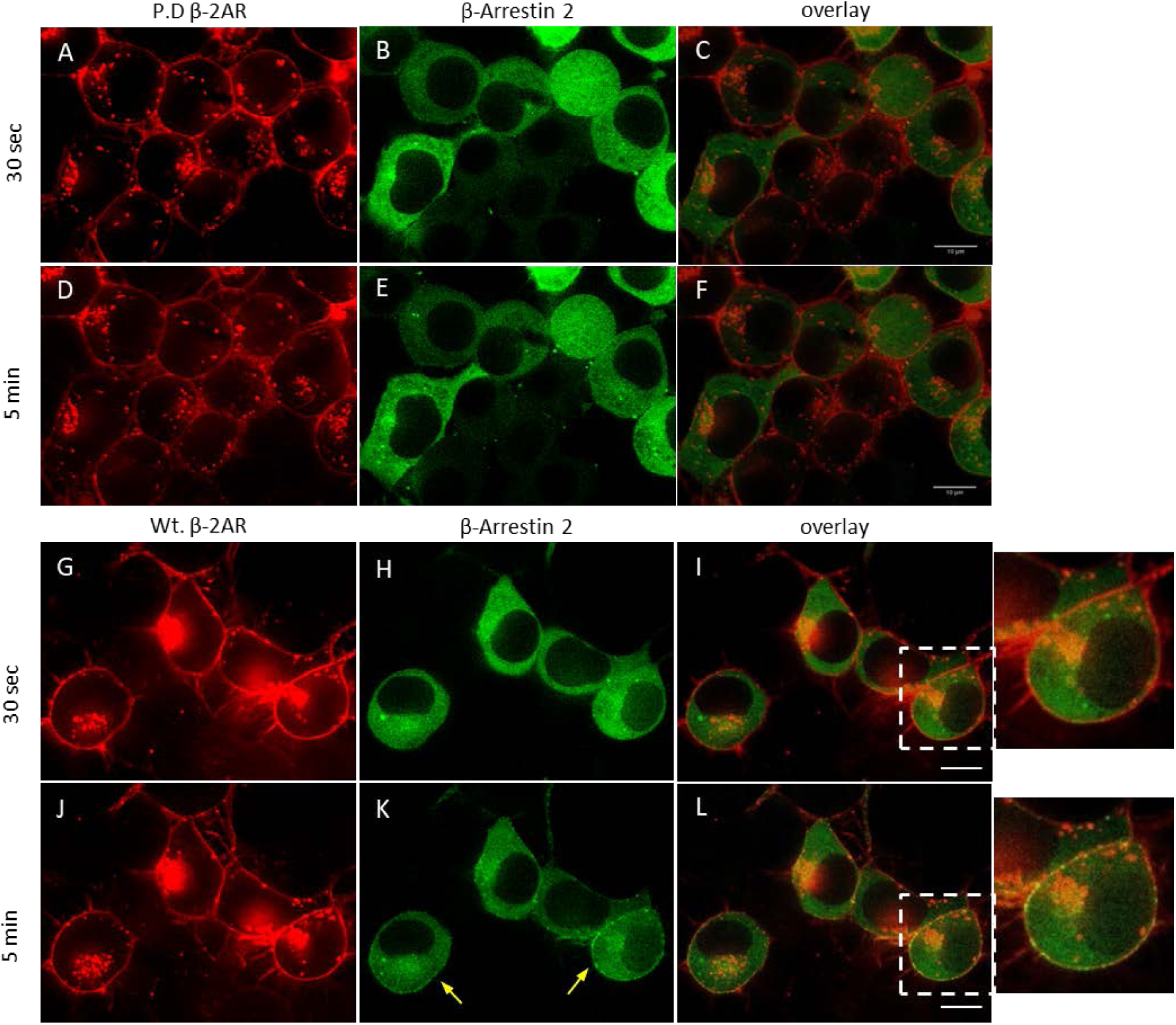
**(A-F)** Spinning disc confocal microscopy images of N2a cells transfected with β_2_AR-P.D.-mCh., arrestin-3-177-mEGFP and wild-type GRK2. **A.** Taken from mCh. channel after 30-second Iso stimulation. **B.** Taken from EGFP channel after 30-second Iso stimulation. **C.** Overlay of **A** and **B**. **D.** Taken from mCh. channel after 5-minute Iso stimulation **E.** Taken from EGFP channel After 5-minute Iso stimulation. **F.** Overlay of **D** and **E**. (**G-L**) Spinning disc confocal microscopy images of N2a cells transfected with β_2_AR-mCh., arrestin-3-177 mEGFP, and wild-type GRK2. Spinning disc confocal microscopy images of N2a cells transfected with β_2_AR-mCh., arrestin-3-177 mEGFP, and GRK2. **G.** Taken from mCh. channel after 30-second Iso stimulation. **H.** Taken from EGFP channel after 30-second Iso stimulation. **I.** Overlay of **G** and **H**. **J.** Taken from mCh. channel after 5-minute Iso stimulation **K.** Taken from EGFP channel After 5-minute Iso stimulation. Yellow arrows show arrestin-3 colocalizations with β_2_AR. **L.** Overlay of **J** and **K**. Magnification 63X, scale bars are 10 µm.

Next, arrestin-3 recruitment to WT and phosphorylation-deficient β_2_AR with or without test compounds or Iso were analyzed by FRET^40^ using membrane-targeted Gap43-mCh.-meGFP as a positive control. After assessing cytotoxicity of L^SH-1^, L^SH-2^, and L^SH-3^ by succinate dehydrogenase activity (MTT) assay^41^ in N2a and HEK293 cells (Supplementary Fig.7), experiments were carried out using 100 µM of each test compound.

To perform FRET experiments in a phospho-deficient system using a plate reader, we validated that the FRET/Donor ratio calculated from the plate reader correlates with % FRET efficiency measured by confocal fluorescence microscopy, analyzed with the ImageJ PixFRET plug-in. N2a cells overexpressing the control FRET construct, Gap43-mCh.-mEGFP, were first imaged by microscopy, resulting in a 10–15% FRET efficiency. Subsequently, the same experiment was conducted using the plate reader, yielding a FRET/Donor ratio of 0.12, which aligned with the microscopy measurements. This consistency is largely due to the minimal bleed-through of these fluorescent proteins into each other’s channels, which was negligible under our imaging conditions (data not shown).

For further experiments, FRET/Donor ratios calculated from plate reader measurements were used. Basal FRET/Donor ratios (prior to stimulation) and response to Iso (20 µM) were measured in N2a cells expressing either wild-type (WT) β2AR or phosphorylation-deficient β2AR together with arrestin-3 (Fig. 5). N2a cells expressing WT β2AR showed approximately 75% higher FRET/Donor signal compared to those expressing the phosphorylation-deficient β2AR (Fig. 5A). Iso treatment further increased the FRET/Donor ratio in the WT system, while no increase was observed in the phospho-deficient system (Fig. 5B). These results indicate that arrestin-3 recruitment to the phosphorylation-deficient β2AR was significantly reduced.

**Figure 5.**
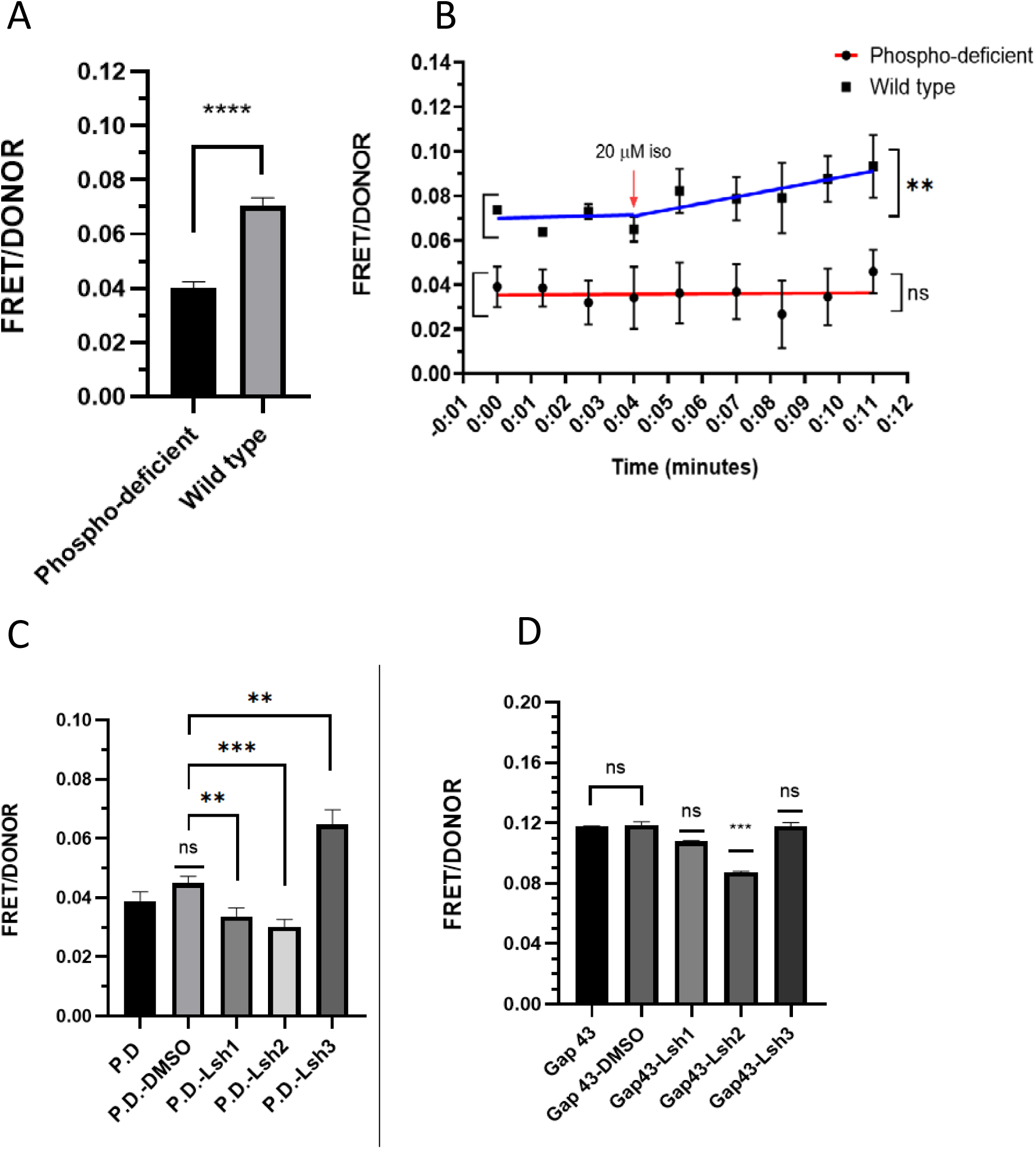
**A.** Basal FRET levels in N2a cell Transfected with β_2_AR phosphorylation-deficient mCherry, arrestin-3-177-mEGFP, GRK2 (negative phosphorylation-deficient.) or β_2_AR - mCherry, arrestin-3-177-mEGFP, GRK2 (positive wt). **B.** Calculated FRET levels in N2a cell before and after 20 µM Iso stimulation. **p<0,005, ****p<0.0001 statistically significant differences. ns: no significant differences. Data Mean ± s.e.m. **C.** L^SH-3^ chemical (1*H*-Indole-3-acetic acid, 6-[[(6-carboxy-3-cyclohexen-1-yl)carbonyl]amino]-α-(4-ethyl-1-piperazinyl) increases FRET between b_2_AR phosphorylation-deficient–mCh., and arrestin-3-177-mEGFP. Each chemical was tested in two separate experiments n=6. **p<0,005, ***p<0,001, statistically significant differences. ns: no significant differences. Data Mean ± s.e.m. **D.** Chemical interaction with membrane targeting sequence fused mCh. and mEGFP transfected N2a cells. ***p<0.001, statistically significant differences, ns: no significant differences. Chemical-treated samples were compared with Gap 43 vehicle (1% DMSO). Data are shown as Mean ± S.E.M.

Next, we used this system to test whether selected compounds promote interaction between phosphorylation-deficient β_2_AR and arrestin-3. 30-minute measurements were recorded following 30-minute incubation (Fig. 5C). FRET/Donor values for phospho-deficient β_2_AR and with and without 1% DMSO (negative control) were calculated as 0.038 and 0.044, respectively. FRET/Donor values for L^SH-1^, L^SH-2^, and L^SH-3^ were found to be 0.033, 0.030, and 0.064, respectively. Compound L^SH-3^ increased FRET by nearly 50%, which suggested that L^SH-3^ facilitates interaction between phosphorylation-deficient β_2_AR and arrestin-3.

To evaluate if the compounds interfered with mEGFP and mCh fluorescent proteins, and therefore impacted measured FRET, another FRET experiment was performed in the presence of test compounds using the positive control Gap43-mCh.-mEGFP. Results indicate that L^SH-1^ and L^SH-3^ do not alter the FRET in this construct, while L^SH-2^ decreases FRET by around 25%. This suggests that the impact of L^SH-1^ and L^SH-3^ on FRET in phosphorylation-deficient β_2_AR-arrestin-3 system is not due to non-specific interference, confirming that L^SH-3^ increased FRET by bringing phosphorylation-deficient β_2_AR and arrestin-3 close to each other.

### L^SH-3^ effect is receptor-specific

We tested the effect of L^SH-3^ on arrestin-3 recruitment to β_2_AR and muscarinic M2 receptor (M2R) using an independent NanoBiT-based assay in HEK293 cells lacking endogenous arrestin-2 and arrestin-3^42^. Cells were co-transfected with SmBiT-arrestin-3 and LgBiT fused at β_2_AR or M2R C-terminus. and pre-treated cell with 100 μM L^SH-3^ (DMSO served as a negative control) for 30 min. In case of β_2_AR, L^SH-3^ treatment significantly enhanced arrestin-3 binding to the receptor in the absence of an agonist (Fig. 6E). Notably, the maximum arrestin-3 recruitment to the isoproterenol-activated β_2_AR remained unchanged, at all isoproterenol concentrations (Fig. 6A,C). In contrast, L^SH-3^ treatment showed no significant effect on arrestin-3 recruitment to M2R, regardless of the presence of agonist (Fig. 6F vs 6B, D). Thus, L^SH-3^ selectively stimulates arrestin-3 recruitment to the β_2_AR, but not to M2R.

**Figure 6.**
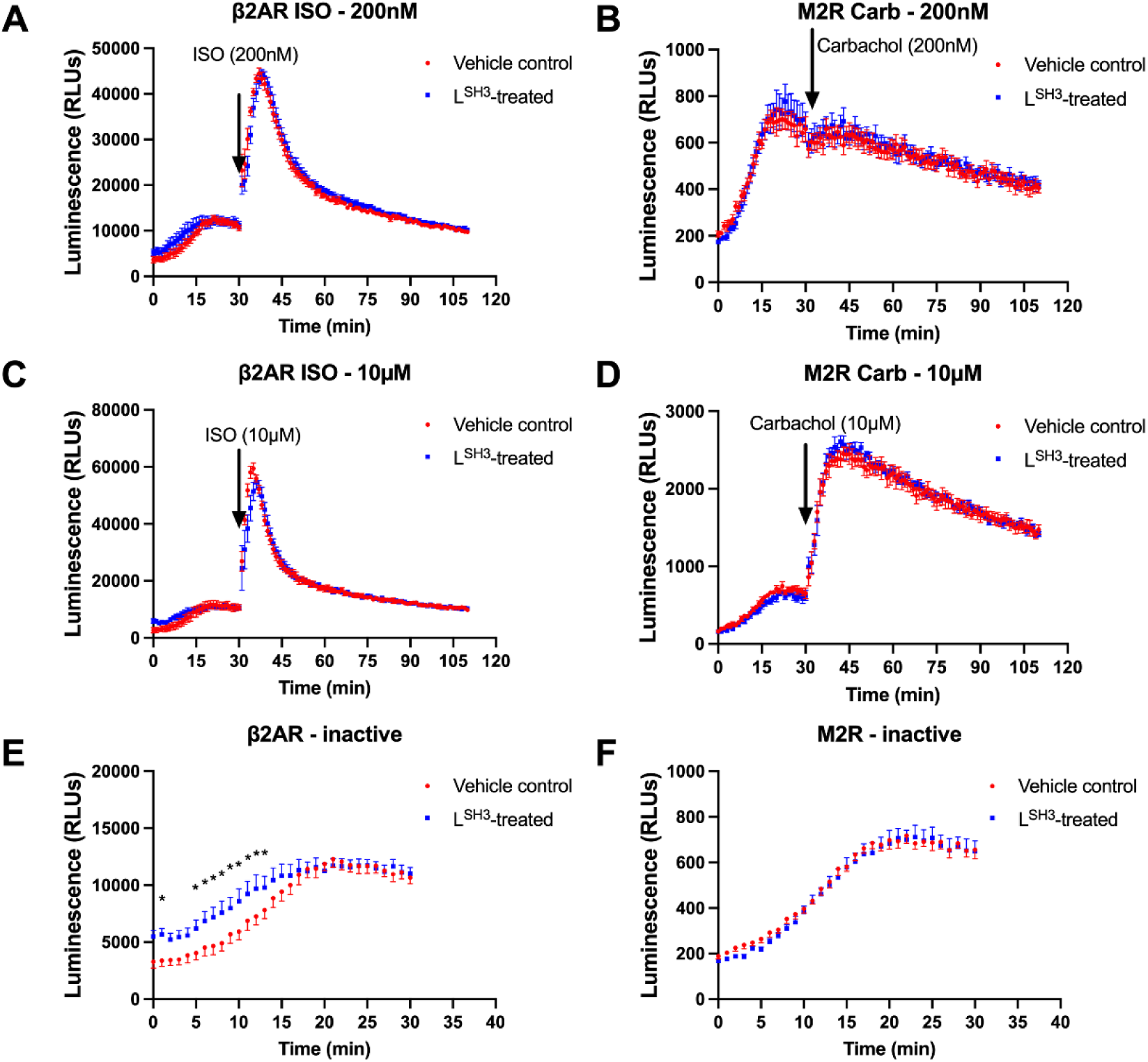
L^SH-3^ treatment selectively enhances arrestin-3 recruitment to inactive β2-adrenergic receptor (β2AR). (**A-D**) NanoBiT assay shows the effect of L^SH-3^ treatment on arrestin-3 recruitment to β2AR and M2R in response to agonist stimulation. HEK293 arrestin2/3 KO cells were co-transfected with SmBiT-arrestin-3 and LgBiT-β2AR or -M2R and treated with 100 μM L^SH-3^ or DMSO (control) for 30 minutes prior to the assay. Agonist-induced arrestin-3 recruitment was measured following stimulation with 200 nM (**A**, **B**) or 10 μM (**C**, **D**) of isoproterenol (β2AR) or carbachol (M2R) at 30 minutes. (**E**, **F**) arrestin-3 recruitment to inactive receptors (0-30 minutes) in the presence or absence of STK treatment. L^SH-3^ significantly increased arrestin-3 binding to inactive β2AR, particularly in the first few minutes (**E**), while no significant effect was observed for M2R (**F**). Data are presented as mean ± S.E.M. (n = 6 for A-D, n = 10 for E-F). Statistical significance was determined by two-way repeated measures ANOVA followed by Fisher’s LSD post hoc test. *p < 0.05.

## Discussion

G protein-coupled receptors are involved in numerous signaling pathways in the body. Precise regulation of initiation and termination of the signal is crucial for the homeostasis of the cell. One of the mechanisms that cells use for terminating the GPCR-mediated signaling is the phosphorylation of certain cytoplasmic receptor residues. Recognition of phosphoresidues by arrestin stabilizes the pre-activated conformation and contributes to GPCR affinity. Dysregulation of phosphorylation prevents arrestin binding and leads to various diseases.^27,43^

Enhanced arrestin mutants were designed to increase arrestin affinity for phosphorylation-deficient GPCRs, but their use requires gene therapy. The effect of phosphorylation in inducing arrestin pre-activation can be recapitulated by phosphopeptides mimicking the C-tail of the receptor^44^, as shown by crystal structures of phosphopeptide-bound arrestin complexes. Other negatively charged molecules like heparin or IP6 mimic the effects of phosphopeptides^45^. However, these compounds cannot be a viable therapeutic option because they do not cross the plasma membrane. We found that the conformational equilibrium of arrestin can be perturbed by small molecules.

We focused on arrestin-3, since available evidence indicates that it is the most conformationally flexible non-visual subtype^31,33^. We confirmed its flexibility by MD simulations and identified a potential binding pocket that emerges upon its transition to pre-activated state. We discovered *in silico* compounds that fit one of these pockets and demonstrated experimentally that one of the predicted compounds, L^SH-3^, binds arrestin-3. In MD simulations L^SH-3^, predicted to stably bind to the back loop, stabilizes its unfolded conformation, and transmits the perturbation from the back loop to the gate loop, triggering its translocation from the polar core towards the lariat loop, facilitating domain inter-domain rotation. Importantly, the back loop was shown to be perturbed upon binding PIP_2_ which led to arrestin-3 pre-activation^24^, and upon arrestin-2 pre-activation^46^. Interestingly, these conformational rearrangements occurred without full dislocation of the C-tail of arrestin-3, which is consistent with our MD simulations, but opposed to the common mechanism of phosphorylation-dependent activation, where the C-tail is displaced upon binding to the phosphoreceptor. Interestingly, it was previously shown that the C-tail of arrestin-3 can be localized close to its N-domain upon binding to the phosphorylated receptor^47^.

We demonstrated experimentally that in the presence of L^SH-3^ arrestin-3 binds phosphorylation-deficient, as well as unstimulated β_2_AR. It should be noted that “unstimulated” does not necessarily refer to the inactive conformation, as β_2_AR samples active conformations even in the absence of an agonist^48^.

Our data suggests that L^SH-3^ acts as a facilitator of arrestin-3 binding to β_2_AR. Also, finding that L^SH-3^ does not enhance arrestin-3 binding to M2 muscarinic receptor suggests that arrestin-3 bound to different GPCRs assumes distinct conformations. Indeed, the back loop is helical in the crystal structure of arrestin-2/M2R complex^18^, while it is unfolded in the arrestin-2/β1-adrenergic receptor complex^17^, suggesting a receptor-specific modulation. It is tempting to speculate that L^SH-3^ might modulate receptor-specific binding of arrestin-3 by stabilizing the back loop in the unfolded state.

The functional effect of the ligand-induced structural modulation of arrestin in cells expressing β_2_AR is currently under study. Our preliminary data in human cardiomyocytes derived from induced pluripotent stem cells^49^, where the β_2_AR /arrestin-3 signaling plays a major role in homeostasis and cardiac dysfunction, suggest an increase in the beating rate in the presence of L^SH-3^. Whether the effect we observe is receptor independent or mediated by β-arrestin2 dependent signaling, similarly to what is observed for some pepducins ^50^, remains to be evaluated and will be the subject of future research.

Overall, our results indicate that arrestin-GPCR interaction can be enhanced by small molecules in a receptor-specific manner. Molecular tools that regulate the interactions of arrestins with GPCRs have significant research and therapeutic potential. Further studies are necessary to explore the uses and limitations of this novel approach.

## Materials and Methods

### Modeling

We used the crystal structure of bovine arrestin-3 (PDB-ID:3P2D, chain B) as a template to model human arrestin-3 (sequence similarity: 96.2%). The missing residues were built using Swiss-Model^51^ while the C-terminal tail was modeled using Modeller software^52^ as it contained a higher number of missing residues than could be handled by Swiss-Model. The C-terminal tail in the model structure, which was provided by the Modeller, was folded back on the N-terminal domain of arrestin-3. Since the C-terminal tail interacts with the rest of arrestin in the basal state and is detached upon activation, we used that lowest-energy model as the initial structure in molecular dynamics simulations. The protonation states of residues were determined using the PROPKA^53^ methodology which computes pKa values of ionizable residues by accounting for the effect of the protein environment at pH=7. The parametrization of small molecules, which were used in simulations of complexes with arrestin-3, was done using CHARMM-GUI^54^ .The output files were examined to check the penalty values associated with parametrization. CHARMM-GUI was also used for solvation and ionization of the systems studied, which was done by neutralizing the systems by 0.15 M KCl. We used CHARMM36m (Chemistry at Harvard Macromolecular Mechanics force field)^55^ and TIP3P^56^ to model protein and water molecules, respectively.

### Molecular dynamics simulations

Atomistic MD simulations were performed on both arrestin-3 and arrestin-3-compound complexes using GROMACS 4.5.0.1^57^ package and the following settings. The particle mesh Ewald (PME)^58^ method was used for the electrostatics with a real space cut-off of 1.0 nm and a grid spacing of 0.13 nm. The Lennard-Jones (LJ) interactions were determined with a twin-range cut-off scheme of 1.0 and 1.4 nm with the long-range interactions updated every 10 steps. The time step was 2 fs. The temperature and pressure were set at 310 K and 1 atm using the Nose-Hoover thermostat^59^ and Parrinello-Rahman barostat^60^, respectively. We performed three replicas for apo arrestin-3 and two for arrestin-3-compound complexes. The simulations were performed for 1 μs.

### Accelerated molecular dynamics simulations

To further check the conformational properties of arrestin-3 we performed accelerated molecular dynamics (aMD) simulations ^61^ using the dual boost option, where independent boost potentials were applied to the dihedral and potential energy of the system, as implemented in NAMD package^62^. We used values of 4455 kcal/mol, 81, -544852 kcal/mol, and -35198 for describing threshold energy for dihedral, dihedral acceleration factor, threshold energy for the potential, and acceleration factor in the dual boost mode, respectively, which were calculated according to the equation in ref.^63^ in aMD simulations. V_avg_dihedral_ and V_avg_potential_ values were obtained from classical MD simulations.

### Clustering trajectories

Trajectories of arrestin-3 were analyzed in terms of the system local and global structural properties: snapshots were selected based on the values of a set of conformational measures: the state of the aromatic core at the interdomain interface, the conformational state of the back loop, the gate loop, and the inter-domain rotation angle (Fig. 2). We measured the distance between Cα atoms of residues F76 and F245 (F75 and F244 in arrestin-2; F79 and Y250 in arrestin-1) for the aromatic core, K313 and G317 (R312 and G316 in arrestin-2; K318 and G322 in arrestin-1) for the back loop, and D291 and H296 (D290 and H295 in arrestin-2; D296 and H301 in arrestin-1) for the gate loop. We compared these values to those measured in crystal structures of arrestin-1, arrestin-2, and arrestin-3 in their inactive and active states((PDB IDs: 7DF9, 2WTR, 1G4M,1G4R, 3GC3, 5W0P, 3GD1, 1SUJ, 1CF1, 4ZRG, 3UGU, 1AYR, 4J2Q, 3UGX, 3P2D, 1ZSH, 1JSY, 4ZWJ, 6U1N, 6TKO, 6K3F, 7JTB, 7F1W). Accordingly, we classified arrestin-3 conformations as intermediate when the back loop and the gate loop took values in the following ranges: 8.5-11.5 Å, 10.5-11.5 Å, respectively. The upper and the lower values were determined such that they could discriminate between different conformations of the back loop: when the distance was smaller than 8.5 Å, the conformation was α-helix, whereas it was coil when it was greater than 9.5 Å, as verified by STRIDE algorithm implemented in VMD^64^. If the values measured did not fall into the intermediate range, they were classified as either inactive or active when they fell below and above the lower and upper limits, respectively.

### Analysis of inter-domain rotation

The interdomain rotation axis was defined as follows: first, we superimposed the N-terminal domain of a set of crystal structures of arrestin-1, arrestin-2, and arrestin-3 in the inactive and active state (the same PDB IDs given above were used), obtaining a pseudo-trajectory which was then subjected to PCA analysis using the *gmx covar* tool of GROMACS^57^. The first PC projection of this pseudo-trajectory was then analyzed with the hingefind algorithm^65^ to identify the rotation axis of the C-domain relative to the N-domain. Finally, the rotation angle of the center of mass of the C-domain around this axis was calculated for every Molecular Dynamics trajectory after superimposing the N-domain using an in-house program adapted from ref. ^66^ . We classified an interdomain rotation angle as active when positive, i.e. rotated counterclockwise relative to the basal state.

### Virtual screening on selected conformations

The *SiteMap* tool^67^ from Schrodinger software was used to identify binding pockets on the selected snapshots. The pocket candidates were evaluated using the following parameters: the number of site points, hydrophobic/hydrophilic, enclosure, exposure, donor/acceptor character of the pocket, volume, and druggability score (Dscore). The selected pockets at the N-C domain interface, referred to as back loop and gate loop, were used to build a pharmacophore, a theoretical model that is a geometrical representation of the chemical functionalities such as hydrophobicity, the capability of acting as hydrogen donors/acceptors, the aromaticity of the target region. Pharmacophore-based virtual screening was carried out on ZINC compound Database using ZincPharmer ^68^.

### Docking

A set of compounds filtered from the virtual screening step was docked to the binding pocket. Glide docking tool^69^ employed for that purpose, following standard preparation protocols. The candidate molecules were docked flexibly to the target site by SP docking where only the trans conformers of amino acids were allowed. We used residues 123-128/306-308/316 and 168-172/291-299 to dock to the back loop, and the gate loop, respectively.

### Reagents

L^SH-1^ compound (Benzamide, 2-chloro-*N*-[3-[(cyclopropylcarbonyl)amino] phenyl]-5-(methylthio)-) was purchased from Enamine, (ZINC ID:31135800), CAS Registry Number: 1386112-75-9. L^SH-2^ compound (1*H*-Pyrazolo[4,3-*c*]pyridine-3-carboxylic acid, 4,5,6,7-tetrahydro-5-[(2-methyl-1*H*-imidazol-5-yl)carbonyl]-1-(3-phenylpropyl)-) was purchased from ChemBridge, (ZINC ID:67490967), CAS Registry Number: 1309319-17-2. L^SH-3^ compound (1*H*-Indole-3-acetic acid, 6-[[(6-carboxy-3-cyclohexen-1-yl)carbonyl]amino]-α-(4-ethyl-1-piperazinyl)-) was purchased from InterBioScreen (ZINC ID:36365956), CAS Registry Number: 1214645-87-0. Lgl1 compound was purchased from Enamine, (ZINC ID:12915185). All the chemicals were dissolved in dimethyl sulfoxide (DMSO) at 10 mM concentration, then aliquoted at 10 µL in PCR tubes, and kept in a -80 °C freezer for further use.

### N2a cell culture

N2a (mouse neuroblastoma) cells were cultured in 44.5% v/v DMEM with L-glutamine (Gibco, REF 41966-029), 44.5% v/v Optimem® reduced serum medium with L-glutamine (Gibco, REF 11058-021), 10% v/v heat-inactivated Fetal Bovine Serum (Biological Industries REF 04-127-1B), 1% v/v Penicillin/streptomycin solution (Biological Industries REF 03-031-1B). HEK293 cells were cultured in the same reagents with DMEM ratio 90%, and no Optimem. Both cell lines were grown at 37°C in a humidified 95% air/5% CO_2_ atmosphere. The cells were routinely subcultured twice a week.

### Molecular cloning

Tagging the arrestin-3 gene from the 177^th^ position was performed by Restriction Free Cloning, which involves two consecutive PCRs. In the first PCR, the mEGFP (monomeric enhanced green fluorescent protein) gene was amplified with primers that carry complementary to mEGFP gene sequences and 25–30 overhanging base pairs from the arrestin-3 sequence from the 177-178^th^ position. The product of this “first” PCR, which carries flanking regions that are homologous to the arrestin-3 gene, was used as a double-stranded “mega primer” in a second PCR to amplify the whole arrestin-3 carrying plasmid. After the second PCR, 1 µL Dpn1 restriction enzyme was added directly into the PCR mix to digest the parental plasmid (arrestin-3). 2 µL of the final mix was utilized for the transformation of *E. coli* XL 1 Blue. Final constructs were verified by Sanger sequencing and subcloned into an empty pcDNA 3.1(-) for eukaryotic expression.

### Plasmid transfection

H9C2 cells were transfected with plasmid DNA (β_2_AR, arrestin-3, and GRK2) 24 hours prior to cardiac differentiation using Lipofectamine 3000 transfection reagent (Invitrogen, L3000001) based on the manufacturer’s instructions. Briefly, 5.4 µl of each P3000 and Lipofectamine 3000 reagents were used for 900 ng of DNA. After 24 h, the medium was replaced with the differentiation medium containing 1% FBS supplemented with 1 µM RA. Following 72-96 h of transfection, H9C2 cardiac cells were incubated in the absence or presence of 100 µM L^SH-3^ for up to 60 mins, stimulated with 1 µM isoproterenol (USP, 35100) to induce βAR signaling^39,70^ and cAMP was measured at indicated time points.

### Confocal fluorescence microscopy

To examine the arrestin-3 colocalization with β_2_AR at the cell membrane, 80,000 N2a cells were seeded on 35 mm glass-bottom petri dishes (In vitro scientific, CA, USA). Twenty-four hours later, cells were transfected with 400 ng each arrestin-3-177-mEGFP, β_2_AR–mCherry (mCh.)/ β_2_AR P.D.-mCh. and GRK2 carrying pcDNA 3.1-/pcDNA 3 plasmids. Confocal microscope imaging was performed 48 hours post-transfection. Before imaging, the media was aspirated, cells were washed with PBS, and 1 mL of PBS was added. Leica DMI 4000 equipped with Andor DsD2 spinning disc confocal microscopy with Leica 63x/1,32 HCX PL APO oil DIC objective was used for imaging. Cells were excited at 480±10 nm emission collected at 525 ± 15 nm (mEGFP channel), excitation at 585±15 nm and emission collected at 630 ± 20 nm (mCherry channel). FRET analysis of Gap43-mCh.-mEGFP was performed using ImageJ pix-FRET plug-in. First, 60,000 N2a cells were seeded in three 35 mm glass-bottom petri dishes. The next day, samples were transfected with 400 ng. Gap43-mCh.-mEGFP, Gap43-mCh, or Gap43-mEGFP carrying plasmids respectively. At 48 hours post-transfection, media was aspirated, cells were washed with 1 mL PBS, and 1 mL PBS was added to each plate before imaging. Donor spectral bleedthrough was calculated from the Gap43-mEGFP transfected sample by exciting the cells from the donor channel, 480 ± 10 nm emission collected at 525 ± 15 nm and FRET channel, excitation at 480 ± 10, and emission at 630 ± 20 nm. Acceptor bleed through calculated from Gap43-mCh. by exciting the cells from the FRET channel, excitation at 480 ± 10 and emission at 630 ± 20 nm, and acceptor channel, excitation at 585 ±15 nm. and emission collected at 630 ±20 nm. FRET sample (Gap43-mCh.-mEGFP) image was taken from all three channels. The images were processed using ImageJ Pix-FRET plug-in that uses an algorithm that calculates the total bleed through pixel by pixel in three stacks of microscope images. The resulting final image gives the %FRET efficiency map.

### FRET measurements

N2a cells (125,000) were seeded on poly D-lysine (PDL) coated 35 mm plastic petri dishes 24 hours prior to transfection. A control plate in each experiment was transfected with an empty vector, and sample plates that were transfected with β_2_AR-P.D-mCh., arrestin-3-177-mEGFP, and GRK2 encoding plasmids (total DNA amounts of the controls and samples were 1200 ng). After 24 h, transfected cells were washed with PBS and then lifted by 140 µL TrypL-E™ Express (Gibco, Cat # 12605028). Cells were resuspended in a 1.8 mL complete medium and counted. 20,000 N2a cells were seeded in each well of PDL-coated F-bottom opaque black 96-well plates (SPL Life Sciences F bottom 96-well immunoplate #31496). After 24 h, cells were washed twice with 200 µL Hanks’ buffered saline solution (HBSS), and the final volume was adjusted to 150 µL with HBSS containing either 100 µM of tested chemical or 1% v/v DMSO (control). Each chemical was also added to each chemical’s blank group. After adding the chemicals, 96-well plates were incubated at 37 °C for 30 minutes in the cell culture incubator and then measured for 30 minutes at the wavelengths as follows; mEGFP; 470/510, FRET; 470/610, mCh.; 570/610 (excitation bandwidth: 15 nm, emission bandwidth: 25 nm) FRET was calculated as the ratio of FRET/Donor. Molecular Device Spectramax id3 was used for measurement.

The %FRET efficiency was found to be around 12% (data not shown). The same construct was used for FRET calculation in the microplate reader. To determine the cell density to use in further experiments, 10,000, 15,000, and 20,000 cells were seeded in each well, and FRET was calculated as FRET intensity/Donor intensity. According to the results, increased cell density increases the intensity of each channel (mEGFP, FRET, and mCh.), but as expected the FRET/Donor ratio was not affected by the cell density. The FRET/Donor ratio was calculated as 0.125, which correlates with the calculated FRET efficiency in a confocal microscope (around 12%).

To assess possible interference between the compounds and mEGFP and mCh fluorescent proteins, N2a cells were transfected with 400 ng of the positive FRET construct, before FRET measurement in a microplate reader. A 96-well plate was incubated for an hour with each compound (100 µM) in a cell culture incubator and then measured by Molecular Devices Spectramax id3 plate reader using the same settings as for the other experiments.

### MTT toxicity assay

To test the toxicity of the compounds, the MTT (3-(4,5-dimethylthiazol-2-yl)-2,5-diphenyl-2H-tetrazolium bromide) assay was used. N2a and HEK293 cells were seeded in clear F-bottom plastic 96 well microplates (Sarstedt AG&Co. KG, Germany, REF: 83.3924) in 100 µL final volume in complete medium, at a density of 10,000/well and 20,000/well for each cell line, respectively, and cultured at 37 °C in 5% Co_2_ incubator for 20 h. The test compounds were added at 50 and 100 µM and incubated for another 4 h. 10 µL MTT solution (5 mg/mL dissolved in PBS) was added to each well and incubated for another 4 h. After that, 100 µL of detergent solution (10% w/v SDS in 0.01 M HCl) was added to each well to dissolve the formed formazan crystals. The plates were shaken for 5 minutes and incubated at room temperature for 16 h. Absorbance was measured at 570 nm with Thermo Scientific Varioskan Lux microplate reader.

### Saturation Transfer Difference (STD) – NMR

For the STD-NMR experiments, arrestin-3 was produced as 6xHis-tagged recombinant protein in *E. coli* using vector pET28a from Twist Bioscience (South San Francisco, California, USA), as described ^71,72^. The L^SH-3^ compound was obtained by InterBioScreen with a non-stereoselective synthesis. It contains three stereocenters and thus can in principle exist as 8 distinct stereoisomers. The commercially acquired compound was first characterized and a set of signals, corresponding to four of the possible diasteroisomers, were identified, with one of them accounting for the 50%. Therefore, to assess the ability of L^SH-3^ to bind arrestin-3, multiple STD spectra were acquired using different compound/protein samples, prepared to better characterize the binding mode. Reported STD data were calculated considering the main isomer.

The protein-compound samples were prepared in a 200:1, 500:1, 1500:1 and 3000:1 compound/protein ratio. The final concentration of the protein was 5 μM, and the final volume was 180 μL. The buffer used is a 20 mM deuterated phosphate buffer pH 7.4. A percentage (less than 5%) of DMSO–*d*6 was added when necessary to improve the solubility of the compound.

^1^H-STD NMR experiments were performed on a 600 MHz Bruker Avance spectrometer. The probe temperature was maintained at 298 K. In the STD experiments, water suppression was achieved by the excitation sculpting pulse sequence.

The on-resonance irradiation of the protein was performed at 0 and 8,8 ppm. Off-resonance irradiation was applied at 40 ppm, where no protein signals are visible. Selective presaturation of the protein was achieved by a train of Gauss-shaped pulses of 50 ms length each. The STD-NMR spectra were acquired with an optimised total length of saturation train of 0.5 s, 1 s or 2 s. Blank experiments were conducted in absence of protein in order to avoid artefacts. The different signal intensities of the individual protons are best analysed from the integral values in the reference and STD spectra, respectively. (I_0_ - I_sat_)/I_0_ is the fractional STD effect, expressing the signal intensity in the STD spectrum (ηSTD) as a fraction of the intensity of an unsaturated reference spectrum. In this equation, I_0_ is the intensity of one signal in the off-resonance or reference NMR spectrum, I_sat_ is the intensity of a signal in the on-resonance NMR spectrum, and I_0_ - I_sat_ represents the intensity of the STD NMR spectrum. The STD NMR binding epitope was calculated based on the normalized STD values from the initial slope of each proton^73^, the initial growth rates approach consists in analyzing the protein–compound association curve using STD values at the limit of zero saturation time, when virtually no compound rebinding or relaxation takes place. The distance between the compound and the protein surface is expressed as absolute STD percentage (STD Abs) and the compound moieties interacting with the macromolecule are illustrated through a color-coded epitope map, obtained by normalizing all measured STD intensities against the most intense signal (to which is assigned an arbitrary value of 100%).

The STD amplification factor (STD-AF) is obtained multiplying ηSTD by the excess of compound and is proportional to the concentration of the protein–compound complex in solution. The curves were built with four concentrations (from 0.3 mM to 5 mM) and the K_D_ values were extracted from the binding isotherm with STD-AF measured at saturation times of 2s. To calculate the STD Amplification Factor (STD-AF), which relates the protons’ proximity to the protein surface, proton H14, H15, and H17 were selected as important moieties for the interaction. The experimental data were fitted with an exponential function.

### NanoBiT assay

NanoBiT measurements were performed using HEK293 cells with CRISPR/Cas9-mediated knockout of arrestin-2 and arrestin-3. Cells were co-transfected in 24-well plates using jetOPTIMUS transfection reagent (Polyplus, Cat. #101000025) with plasmids encoding small-bit (SmBiT)-tagged arrestin-3 and large-bit (LgBiT)-tagged β_2_AR or muscarinic M_2_ receptor (M_2_R). At 24 hours post-transfection, cells were transferred to 96-well white-walled plates (30, 000 cells/well in 100 μL serum-free DMEM) and serum-starved for 16 hours. At 48 hours post-transfection, cells were pre-treated with either L^SH3^ (100 μM) or DMSO (control) for 30 minutes at 37°C in a humidified atmosphere containing 5% CO_2_. The nanoBiT substrate (Nano-Glo, Promega, Cat. #N2011) was prepared according to manufacturer’s instruction and added (20 μL/well). Luminescence was monitored at 1-minute intervals for 30 minutes using BioTek Synergy Neo2 microplate reader until signal stabilization. Following equilibration, receptor-specific agonists were added (5 μL/well) to a final concentration indicated (200nM or 10μM): isoproterenol for β_2_AR-expressing cells or carbachol for M_2_R-expressing cells. Luminescence measurements continued at 1-minute intervals for an additional 80 minutes.

### Statistical analysis

All experiments were performed in triplicates, and the results were presented as mean ±S.E.M (Graphpad Prism version 8.0.1). Statical analyses were carried out in GraphPad Prism. For a single experiment, two-way ANOVA followed by the Tukey test was used, to analyze two independent experiments Nested t-test was used. Line graphs were tested with the Pearson r correlation test.

## Acknowledgements

This study was funded by the Scientific Research Council of Turkey (TUBITAK), project number 117Z245. β_2_AR - mCh and β_2_AR -phosphorylation-deficient – mCh. and Gap43-mCh.-mEGFP carrying plasmids were provided by former Son’s Lab member, Hüseyin Evci, and arrestin-3 carrying plasmid was a kind gift from Mark Scott (CNRS Research Associate, Paris, France). GM gratefully acknowledges the help of Dr. Zhenlong Li in the Harel Weinstein Lab, Weill Cornell Medicine, who provided the starting full-length molecular model for arrestin-3; she also acknowledges funding from Spoke 7 of Programma di ricerca CN00000013 “National Centre for HPC, Big Data and Quantum Computing” of the NextGenerationEU initiative. The work at Vanderbilt University was funded by NIH grants EY011500 and GM122491.

## Supplementary Information

**Supplementary Fig. 1.**
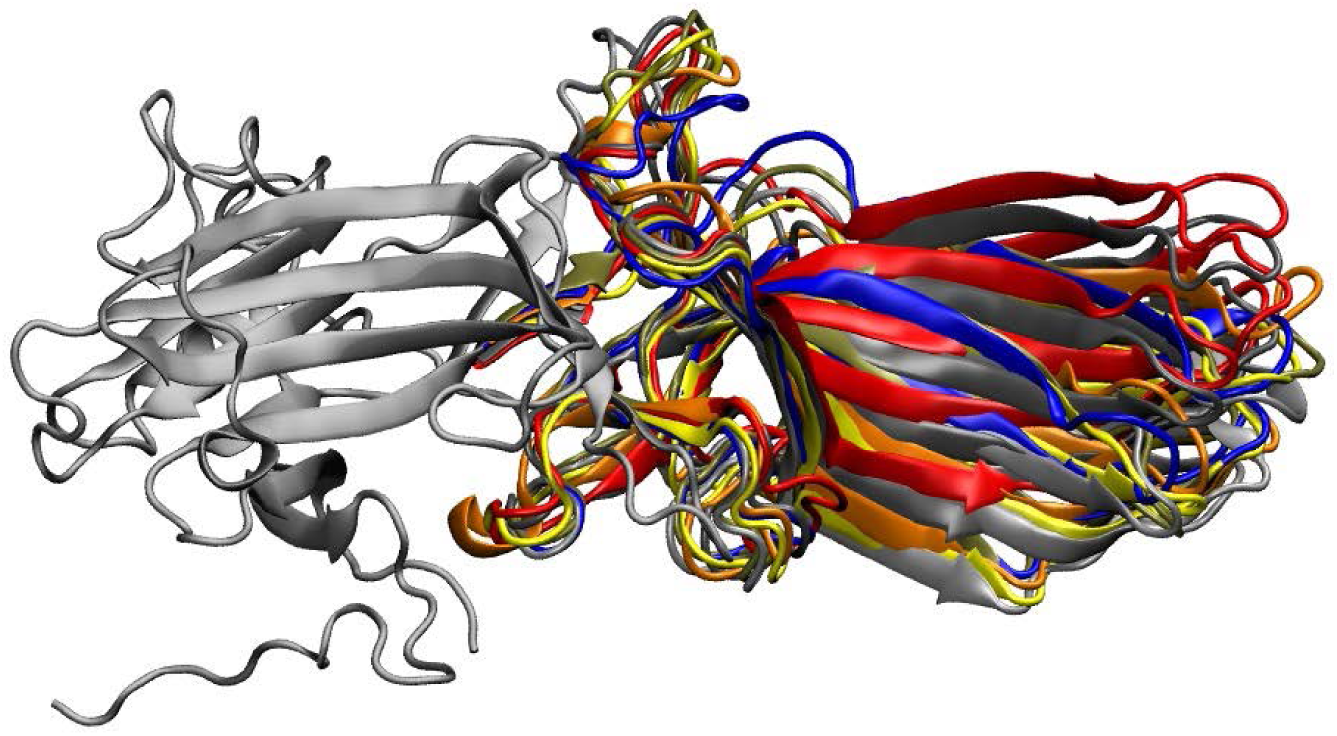
Selected structures from Arr3 MD trajectories. The structures are aligned with respect to the N domain of the protein which is shown in gray. The C domains of different structures are shown in different colors.

**Supplementary Table 1.** Structural parameters used to describe the conformational states of *short helix* and *gate loop* as well as inter-domain rotation angle.

| Snapshot # | Short helix (SH) (Å) | Gate Loop (GL) (Å) | Conformational States of SH and GL | Rotation angle (°) |
| --- | --- | --- | --- | --- |
| 1 | 4.2 | 10.5 | Inactive, Intermediate | -0.5 |
| 2 | 6.9 | 10.5 | Inactive, Intermediate | 9.8 |
| 3 | 8.5 | 8.3 | Intermediate, Inactive | 9.6 |
| 4 | 5.9 | 11.2 | Inactive, Intermediate | -10 |
| 5 | 10.3 | 9.9 | Intermediate, Inactive | -6.7 |
| 6 | 11.5 | 10.5 | Intermediate, Intermediate | -6.3 |
| 7 | 10.9 | 10.6 | Intermediate, Intermediate | -6.2 |

**Supplementary Table 2.** Druggability scores and ranking of selected pockets.

| Snapshot# | Dscore | Sitemap ranking |
| --- | --- | --- |
| 1 (gate loop) | 1.06 | 1/10 |
| 2 (Short helix) | 0.95 | 1/10 |
| 3(Short helix) | 0.95 | 1/10 |
| 4(Short helix) | 0.92 | 1/10 |
| 5(Short helix) | 1.03 | 2/10 |
| 6(Short helix) | 1.04 | 1/10 |
| 7(Short helix) | 0.95 | 2/10 |

**Supplementary Table 3.** Glide binding scores of the compounds that target *short helix*.

| ZINC ID | Binding_Score (kcal/mol) |
| --- | --- |
| 162 | -8.91 |
| 11867617 | -8.73 |
| 6812752 | -8.22 |
| 67490967 | -8.38 |
| 36075880 | -7.97 |
| 2560972 | -7.71 |
| 14971552 | -7.20 |
| 31135800 | -8.58 |
| 16925920 | -9.84 |
| 36365956 | -8.27 |

**Supplementary Fig. 2.**
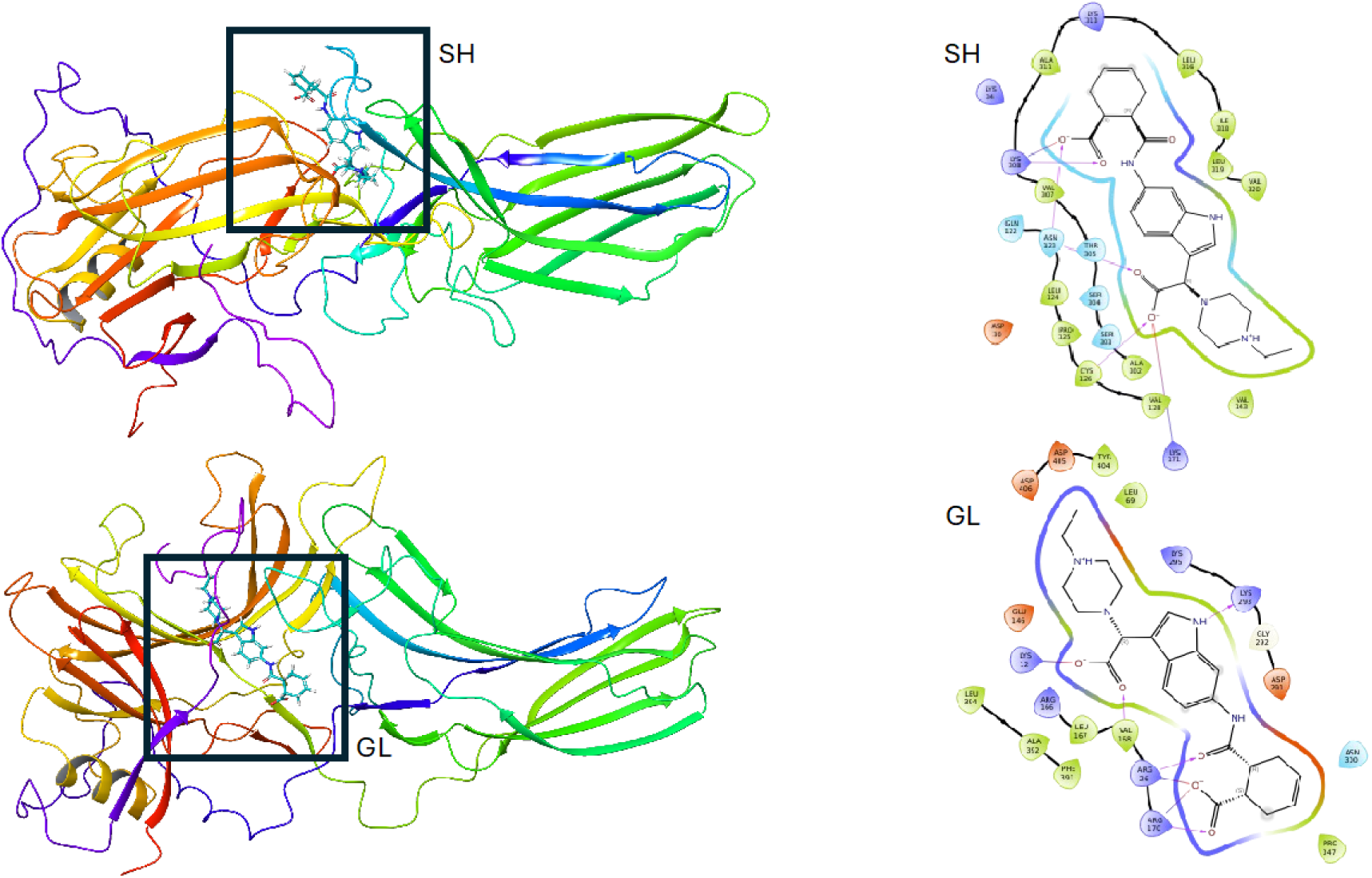
Binding poses of the compound L^SH-3^ docked at the *short helix (SH)* and the *gate loop (GL)* binding pocket.

**Supplementary Fig. 3.**
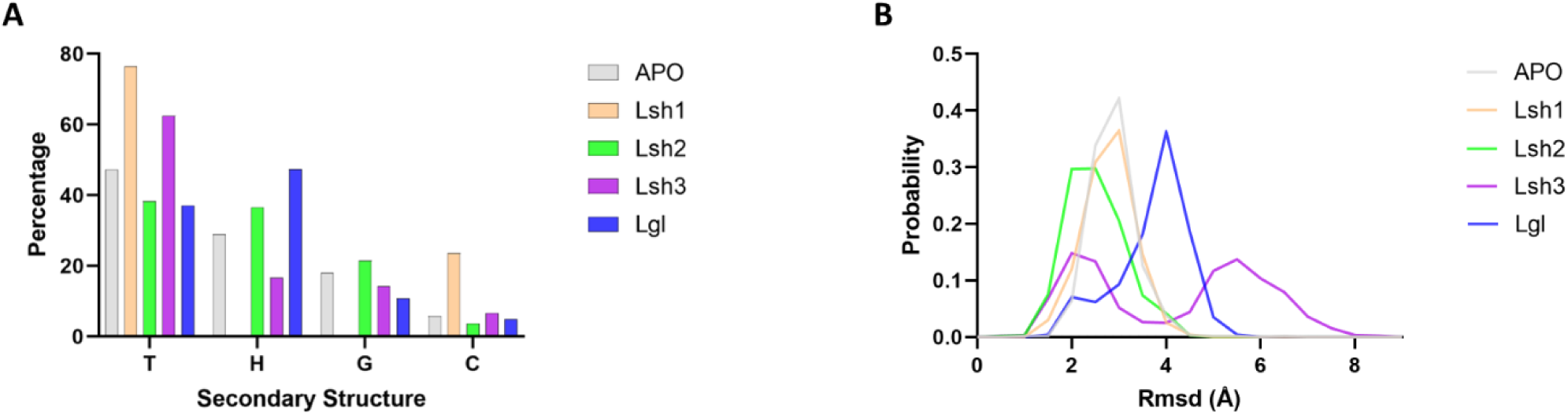
Percentage secondary structure content of the *short helix* sampled in *apo* and ligand-bound Arr3 trajectories calculated by ‘DSSP’ algorithm implemented in VMD, T, H, G, and C correspond to turn, α-helix, 310-helix, and coil. **B.** The RMSD of the *gate loop* calculated using Cα atoms of residues 291-299.

### NMR characterization

**Supplementary Table 4.**
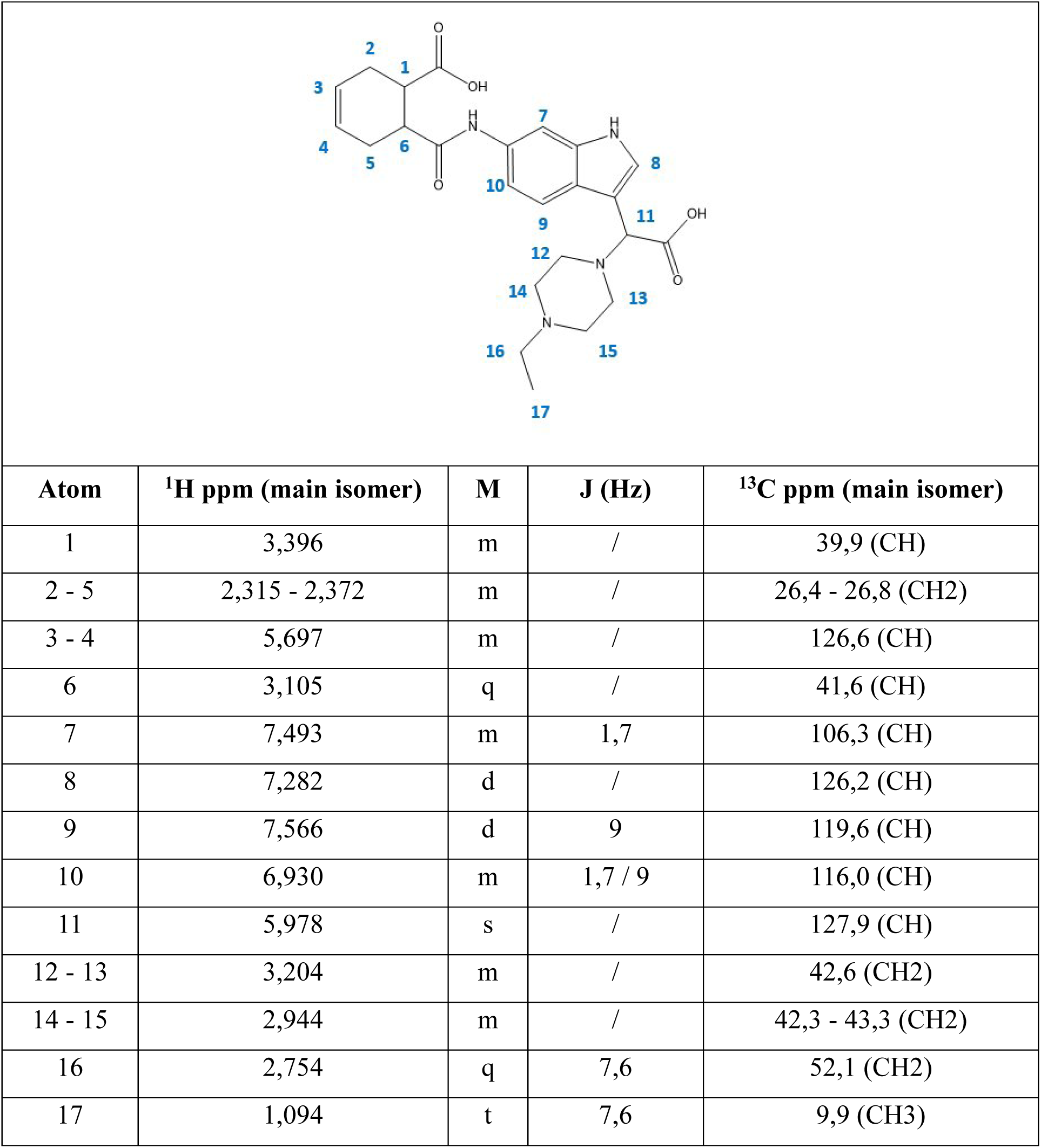
Structure, ^1^H chemical shifts, multiplicity, J values and ^13^C assignments of L^SH-3^ obtained in phosphate buffer at 298K.

|  |  |  |  |  |
| --- | --- | --- | --- | --- |
| Atom | $^1\text{H}$ ppm (main isomer) | M | J (Hz) | $^{13}\text{C}$ ppm (main isomer) |
| 1 | 3,396 | m | / | 39,9 (CH) |
| 2 - 5 | 2,315 - 2,372 | m | / | 26,4 - 26,8 (CH<sub>2</sub>) |
| 3 - 4 | 5,697 | m | / | 126,6 (CH) |
| 6 | 3,105 | q | / | 41,6 (CH) |
| 7 | 7,493 | m | 1,7 | 106,3 (CH) |
| 8 | 7,282 | d | / | 126,2 (CH) |
| 9 | 7,566 | d | 9 | 119,6 (CH) |
| 10 | 6,930 | m | 1,7 / 9 | 116,0 (CH) |
| 11 | 5,978 | s | / | 127,9 (CH) |
| 12 - 13 | 3,204 | m | / | 42,6 (CH<sub>2</sub>) |
| 14 - 15 | 2,944 | m | / | 42,3 - 43,3 (CH<sub>2</sub>) |
| 16 | 2,754 | q | 7,6 | 52,1 (CH<sub>2</sub>) |
| 17 | 1,094 | t | 7,6 | 9,9 (CH<sub>3</sub>) |

#### STD-NMR

STD experiment can map protons close in space to the protein. STD spectrum is obtained by the difference of two spectra: *on resonance* and *off* resonance, respectively. In the *on resonance* experiment the protein receives a selective irradiation (at frequencies able to excite only the protons of the protein and not affecting the atoms of the ligand) that saturates the macromolecule protons; then due to intramolecular nuclear Overhauser effect (NOE) the magnetization is transferred to the ligand, that is near the protein (4-5 Å), which in turn, by exchange, is moved into solution where it is detected. The *off-resonance* spectrum is acquired with a saturation frequency where no protein or ligand signals are detectable, usually around ∼40 ppm. This off-resonance spectrum represents a normal 1H NMR. Subtraction of the on-resonance and off-resonance spectrum results in the final saturation transfer difference (STD) NMR 1D spectrum showing only signals from ligands that have binding affinity, while the peaks of the distant protons are absent. The analysis of different STD spectra acquired irradiating different frequencies (corresponding to different residues of the protein) allows the identification of the type of protein residues contacting the ligand and gives the possibility to identify the protein binding pocket (see ref.^34^ in main text)

**Supplementary Fig. 4.**
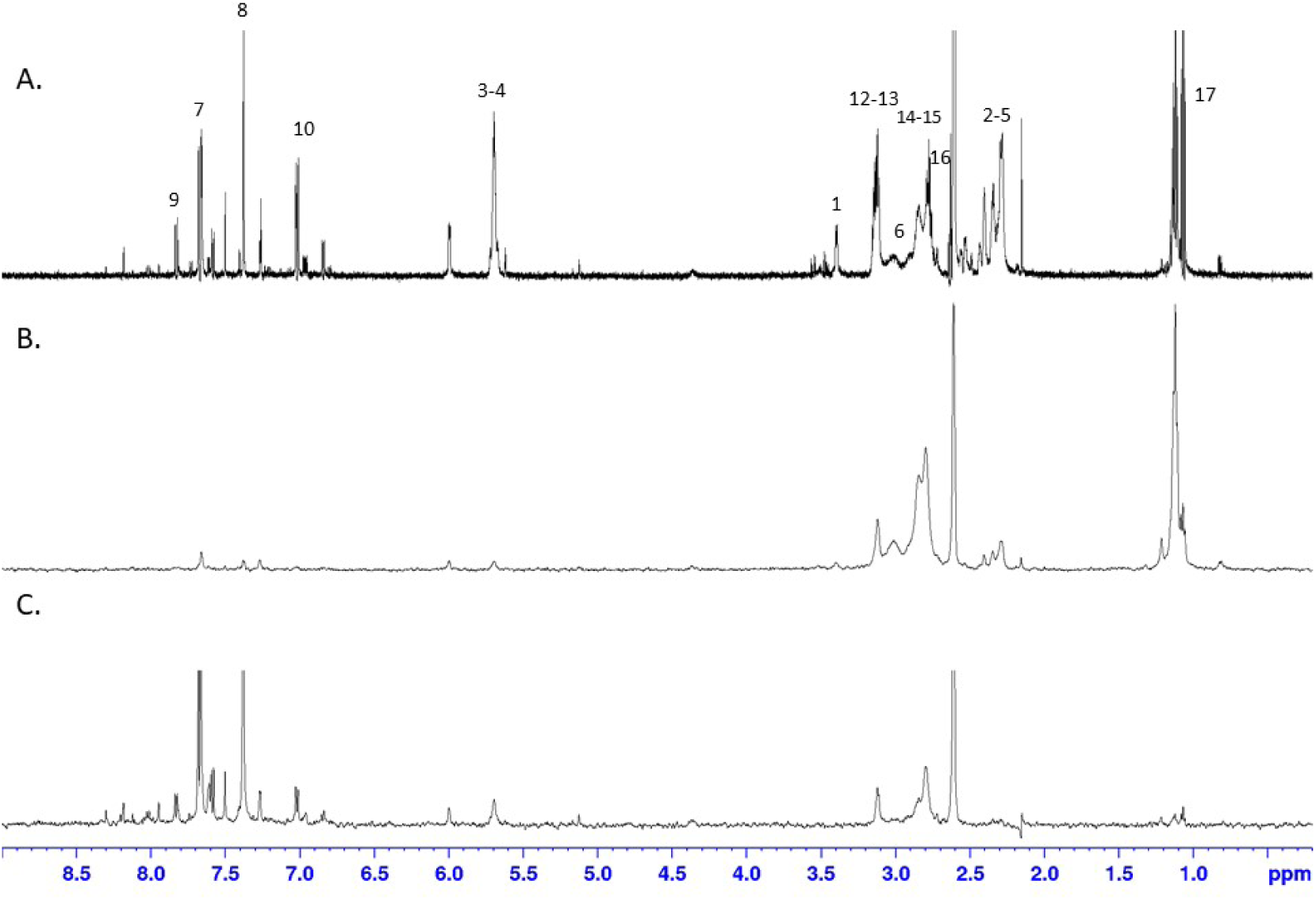
**A.** ^1^H-NMR spectrum of L^SH-3^ **B.** STD spectrum irradiating at 0 ppm **C.** STD spectrum irradiating at 8 ppm

**Supplementary Table 5.** STD Abs at 0 and 8 ppm; the total ligand saturation values are highlighted in the last row.

| H | STDAbs at 0 ppm | STDAbs at 8 ppm |
| --- | --- | --- |
| 2 - 5 | 0,93 | 0,00 |
| 3 - 4 | 0,50 | 0,55 |
| 6 | 5,26 | 0,00 |
| 7 | 1,19 | 3,70 |
| 8 | 0,83 | 1,89 |
| 9 | 1,00 | 3,13 |
| 10 | 1,19 | 1,89 |
| 11 | 0,59 | 1,11 |
| 12 - 13 | 2,00 | 0,55 |
| 14 - 15 | 7,14 | 1,32 |
| 17 | 2,22 | 0,36 |
| **SUM** | **22,86** | **14,50** |

STD was also employed as a fast method for determination of K_D_ values for weakly binding ligands. To get protein-ligand association curves from STD NMR experiments, Mayer and Meyer introduced the conversion from observed experimental intensities to STD amplification factors (STD-AF) by multiplying the observed STD by the molar excess of ligand over protein (Meyer, B.; Peters, T. Angew. Chem. 2003, 42, 864–890).

**Supplementary Table 6.** STD (%) obtained with saturation time of 2s at irradiation frequency 0,0 ppm, 298 K.

| Atom | STD Abs | STD-AF | STD-AF<sub>0</sub> |
| --- | --- | --- | --- |
| 2-5 | 1,07 | 2145,92 | 305,87 |
| 3-4 | 0,29 | 588,24 | 191,17 |
| 7 | 0,50 | 995,02 | 180,52 |
| 8 | 1,41 | 2816,90 | 133,66 |
| 9 | 0,83 | 1666,67 | 299,77 |
| 11 | 1,00 | 2000,00 | 174,24 |
| 12-13 | 2,00 | 4000,00 | 413,80 |
| 14-15 | 6,67 | 13333,33 | 1982,58 |
| 17 | 6,25 | 12500,00 | 731,79 |

In the following figure, the STD-AF values of selected protons of L^SH-3^ are reported as a function of the saturation time. For each proton, STD-AF values were obtained at different saturation times (0.5, 1 and 2 s) and fit by using the equation STD-AF(*t*_sat_)=STD-AF_max_[1−exp(−*k*_sat_*t*_sat_)]. The initial slopes, STD-AF_0_, are obtained from 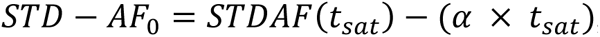, where α is the slope of the curve.

**Supplementary Fig. 5.**
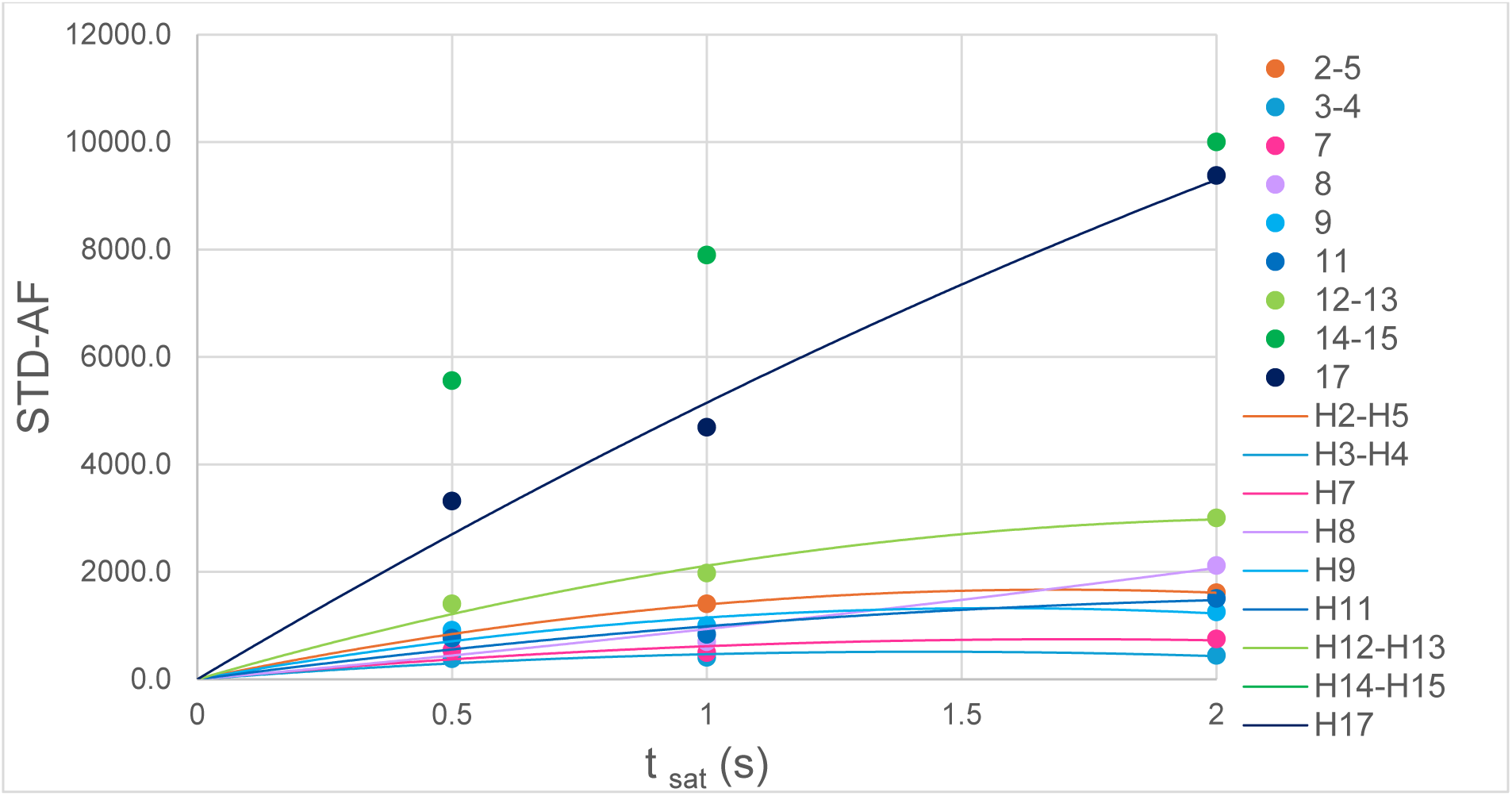
For each proton, STD-AF values were obtained and plotted at different saturation times (0.5, 1 and 2 s). The different coloured dots represent the experimental values and the lines are the mathematical fit to exponential function.

Therefore, a plot of STD-AF values at increasing ligand concentrations will give rise to the protein-ligand binding isotherm, from which the dissociation constant K_D_ can be derived (the K_D_ coincides with the concentration of ligand necessary to reach a 50% of the fraction of bound protein).

**Supplementary Fig. 6.**
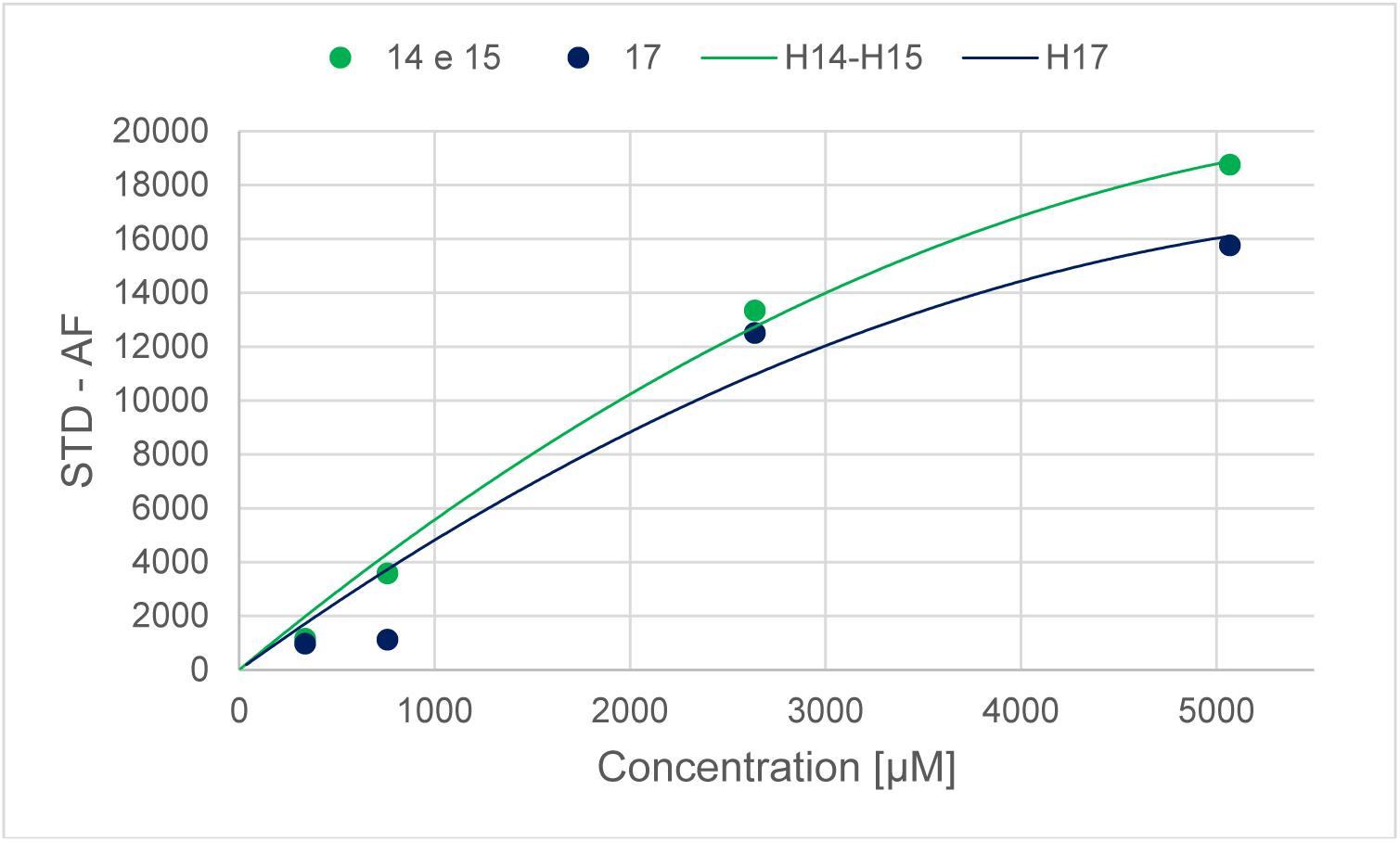
Binding curve of L^SH-3^ with Arr3. The dots represent the experimental values, while the lines are the mathematical fit to exponential function.

#### FRET

##### Cytotoxicity in N2a and HEK293 cells

We assessed the toxicity of L^SH-1^, L^SH-2^, and L^SH-3^ by succinate dehydrogenase activity (MTT) assay^35^ . Each compound was tested at 50 and 100 µM to determine the maximum non-toxic concentration that can be used (1% DMSO served as control). The toxicity of the compounds was determined using Neuro 2A (N2a) cell line, which is a mouse neural crest-derived cell line that has been extensively used to study neuronal differentiation, axonal growth, and GPCR signaling^36^ . Tested compounds were not toxic. No toxicity was either observed in HEK293 cells. Given that some chemicals, such as flavonoids and phytochemicals, can reduce MTT reagent by non-enzymatic reactions^37,38^, we performed an MTT assay in the absence of N2a or HEK293 cells. Each chemical was added to the sample and compared against a solution that contained 1% DMSO. As a result, we did not observe such interference. From these observations, we carried out further experiments using 100 µM of each test compound.

**Supplementary Fig. 7.**
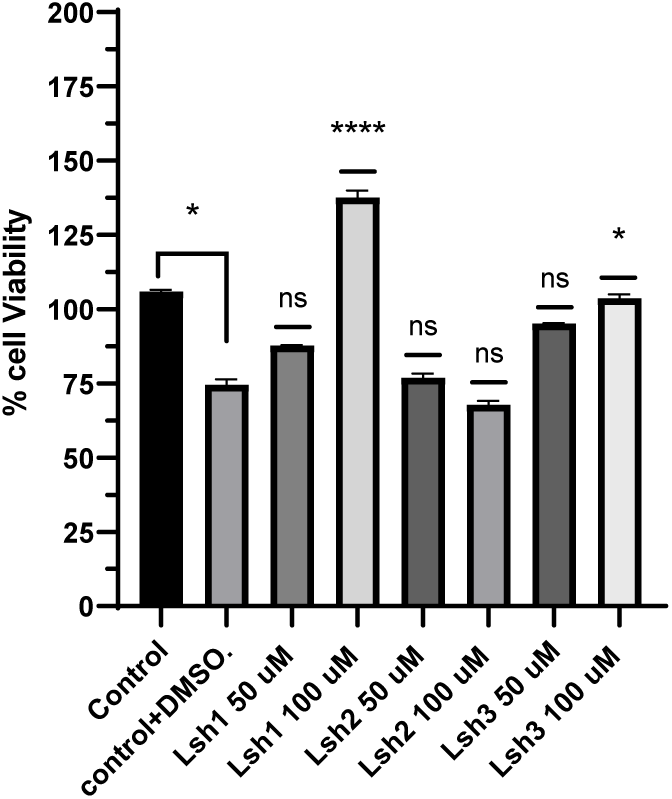
Toxicity test of the compounds in N2a cells. The final DMSO concentration is 1% in all groups except the control group. *p<0.05, ****p<0.0001 statistically significant differences. ns: no significant differences. Data Mean ± s.e.m

## Footnotes

* We use systematic names of arrestin proteins, where the number after the dash indicates the order of cloning: arrestin-1 (historic names S-antigen, 48 kDa protein, visual or rod arrestin; SAG in HUGO database), arrestin-2 (β-arrestin or β-arrestin1; ARRB1 in HUGO database), arrestin-3 (β-arrestin2 or hTHY-ARRX; ARRB2 in HUGO database), and arrestin-4 (cone or X-arrestin; ARR3 in HUGO database).

